# Parallel evolution of full-length genomes in a long-term evolution experiment with phage φX174

**DOI:** 10.1101/2025.04.22.649699

**Authors:** Eva Bons, Hélène Chabas, Hannelore MacDonald, Alejandro Escalera Ledermann, Jenny Dunstan, Nicolas Ochsner, Daniel C. Angst, Sebastian Bonhoeffer, Roland R Regoes

## Abstract

The study of evolution in organisms with high mutation rates requires detailed data on the genomic composition of the population. Here we report on an innovative use of high-throughput sequencing technology combined with rapid experimental evolution that allowed us to follow the genetic diversification of four independent bacteriophage populations that we evolved without actively imposing any selection pressures (in addition to those inherent in the bacterial cell environment) for 412 generations in unprecedented detail. Tracing over 80’000 bacteriophage genomes, representing 884 distinct genotypes, we find that the patterns of viral diversification result in largely non-congruent genotype distributions, but also document multiple instances of parallel evolution. Unlike previous observations of parallelism in viral evolution, we do not just see the parallel evolution of mutations at single sites, but on the level of full-length genomes. Computer simulations that recapitulate our experiments in great detail show that this degree of parallelism is inconsistent with neutral evolution. We further show that the observed extent of parallel evolution biases phylodynamic analysis of migration rates, and wrongly estimates significant migration between the independent evolution lines. Our approach is applicable to many other viruses and paves the way to advanced population genetic investigations that aim to shed light on phenomena that relate to genetic linkage across the genome.

## Introduction

Evolution is driven by random mutations and subsequent changes in the frequency of organisms due to selection and drift. For complex organisms with low mutation rates these two processes typically occur on separate time-scales. For viruses with high mutation rates, however, the generation of new variants and their frequency changes are intertwined, leading to many concurrent variants (Gerrish and Lenski, 1998, Desai and Fisher, 2007), which makes it particularly difficult to disentangle the contribution of the two processes to viral diversification.

The specific population genetic regime that characterizes viral evolution — high mutation rates, and thus concurring variants, and very strong selection pressures — poses particular challenges for population genetic inference schemes that aim to estimate population genetic and dynamical parameters from sequence data, such as phylodynamics. In a nutshell, phylodynamics aims to estimate population biological parameters from sequence data, by first reconstructing phylogenies, then re-interpreting these phylogenies as genealogies, and then estimating key parameters, such as birth and death rates or migration rates between spatial locations, using likelihoods that connect the population biology to genealogies. These inference schemes are usually tested on data that were simulated according to the assumptions of the inference models. In particular, when testing these schemes, selection pressures and their influence on reconstructing genealogies are not taken into consideration. Most importantly, phylodynamics inference methods have not yet been experimentally assessed. To this end, we need to generate data that allow us to trace the evolutionary dynamics of viruses in fine detail.

One technical issue that hampers the fine-grained tracing of the molecular evolution of viruses through time is the lack of sufficiently resolved data on the genomic composition of viral populations. On the one hand, full-genome sequences can be generated by picking single strains from a diverse population and sequencing them. This, however, detects at most a few dozen variants that have mostly high frequency in the population (Keele et al., 2008, Li et al., 2010). On the other hand, early high-throughput sequencing technology could detect low frequency mutations, but generated short reads and thus did not provide linkage information (Acevedo et al., 2014), requiring computational reconstruction of the haplotypes (Zagordi et al., 2011, Prosperi and Salemi, 2012) which is confounded by the strong assumptions that such procedures have to adopt. Here, we report on an approach involving long-read sequencing in combination with computer simulations that overcomes these issues, and successfully determine the compositional changes of four independently evolving populations of φX174, a bacteriophage, during a long-term evolutionary experiment. Our approach results in what we call *whole-population whole-genome sequencing data*.

We chose the bacteriophage φX174 for this proof-of-concept study on whole-population whole-genome tracking of viral evolution because of its short, well-annotated genome of 5.3kB (Sanger et al., 1977), its fast replication cycle of approximately 26 minutes (Baker et al., 2016), and its high mutation rate (Cuevas et al., 2009). For these reasons, φX174 has been one of the bacteriophages of choice in evolution experiments for over two decades (Bull et al., 1997, Wichman et al., 1999, Miller et al., 2016).

The whole-population whole-genome sequencing data we generated gives us a picture of the evolutionary dynamics of φX174 of unprecedented detail, allowing us to trace the emergence and frequency changes of genotypes of the phage, comprehensively down to less than 1%, through time without loosing linkage information. This enabled us to document parallel evolution not just at single loci in the genomes, but of combinations of mutations and even entire genotypes. With these data, we could also assess phylodynamics inference methods and establish systematic biases of these methods when applied to our experimental sequence data.

## Results

### Generating data on the evolutionary dynamics with long-term evolution experiments in a liquid handling platform

To generate data on the evolutionary dynamics, we evolved φX174 in 4 replicates in a constant and controlled environment of its host cells — *Escherichia coli C* (see Figure 1 and Materials and Methods). During the evolution experiment, we did not intentionally impose any selection pressures, for example, by providing unusual host cells to which the phage was not adapted to, or by inhibiting phage replication. We could not rule out that selection pressures were operating though, and provide evidence for selection below. The rationale for not intentionally applying selection was to obtain insights into near neutral bacteriophage diversification and to generate a data set for phylodynamic analysis that conforms to the underlying assumptions of phylodynamics. We seeded each evolution line with the same, genetically heterogeneous stock population of φX174. We then serially transferred 2% of the bacteriophage population into wells containing fresh host cells for 412 generations. We transferred approximately every 26 minutes, consistent with the duration of one replication cycle of φX174. With a newly developed experimental protocol (see Materials and Methods) we then extracted the DNA of 5% of the final bacteriophage population (generation 412), of an intermediate population (generation 196), as well as the original, “ancestor” bacteriophage population (generation 0), and sequenced this DNA using PacBio sequencing (Eid et al., 2009).

**Figure 1:**
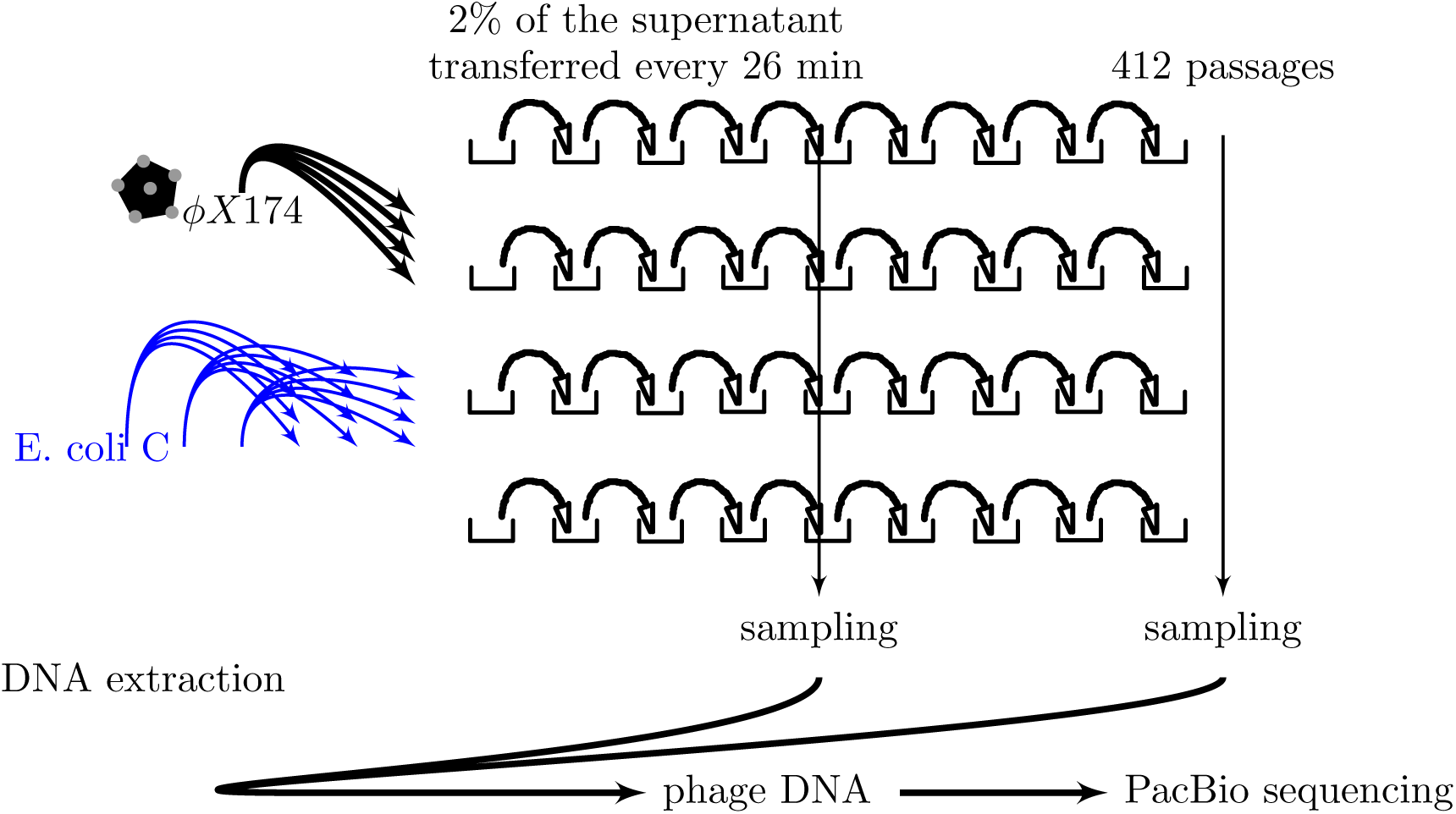
Experimental design. Using a liquid handling platform, we seeded wells containing rich medium with *E. coli C* — the host of φX174. After a delay of approximately 90 minutes that allows the host cells to enter the exponential growth phase, the first column of wells were inoculated with φX174. Subsequently, 2% of the supernatant was transferred every 26 minutes into wells of the next column that had been pre-seeded with host cells. The host bacteria in the wells were always seeded from the stock population, while bacteriophages (along with a few bacteria) were passaged from well to well. Due to the seeding with stock bacteria and the short time between transfers of 26 minutes, only the bacteriophage populations could evolve during the experiment (see Figure S1. After passages 196 and 412, samples were taken, the bacteriophage DNA was extracted, and submitted to PacBio sequencing. For more details see Materials and Methods.

PacBio sequencing requires double-stranded DNA that is turned into a circular template. Because the average read length of PacBio is several times longer than the genome of φX174 — over 38 kb in our experiment — the majority of reads contain multiple, (re-)sequenced copies of each sampled phage genome. By considering the consistency with which each nucleotide has been called at a given position, one can derive a quality score for each position, and generate the so-called circular consensus sequence (Acevedo et al., 2014, Wenger et al., 2019). Thus, the high error rate of PacBio, and other long-read sequencing platforms, of a single nucleotide read (approximately 10-15%) is substantially reduced by reading each position multiple times (Wenger et al., 2019). To further increase the accuracy of our whole-genome sequences, we restrict the circular consensus sequences to quality scores higher than 47, which translates to a sequencing error rate of 2 × 10^−5^ per nucleotide, or to less than one mis-called nucleotide per genome.

After these procedures to ensure sequence quality, we obtain over 110’000 reads covering the entire genome of φX174, representing over 31’000 distinct genotypes (see Figure 2). The majority of these genotypes are only present at very low frequency though. This raises the question whether they were present in the sample or whether they arose due to mutations introduced during the PCR or cell amplification involved in extracting the DNA and preparing it for sequencing, or due to errors introduced during PacBio sequencing that were not corrected through the circular consensus sequences.

### Simulations guide thresholds to exclude sequences generated by DNA extraction or sequencing errors

In order to quantify the extent to which mutations are introduced during the DNA extraction and preparation, we performed *in silico* simulations. These simulations captured the population genetics of the serial passaging experiment, as well as the the methodological steps involved in sampling, preparing and sequencing the viral populations. The exact steps of the experimental setup incorporated in the simulations are shown in Figure 3A and are explained in detail in the Materials and Methods. In brief, the serial passaging experiment is simulated using a previously described sequence evolution simulation tool (Bons and Regoes, 2018), which includes the replication and genetic diversification of the viral population as well as the bottlenecking involved in passaging and has been adapted to φX174. We also incorporated fitness effects of mutations in φX174 that have been experimentally estimated by Domingo-Calap et al. (2009). The simulations also accounted for the methodological steps involved in sampling, preparing and sequencing the viral populations. The sampling of the viral population and the DNA extraction are implemented as a series of bottlenecking and amplification steps that are involved in the protocol. Bottlenecks can lead to the loss of genotypes, while amplification steps can introduce mutations and distort the relative frequencies of genotypes in the sample. Lastly, the simulations incorporate the introduction of sequencing errors at a rate of 2 × 10^−5^ per nucleotide (consistent with restricting the sequencing data to reads with a PHRED score of 47 or higher). The detailed simulation allows us to understand in which exact step sequence variation is introduced, and to which extent this contributes to the variation observed in the final reads.

The simulations allow us to estimate how frequently mutations introduced during the DNA extraction protocol generate genotypes that are not present in the evolved populations. We find that such genotypes with errors do not reach high frequency. Most of the genotypes with errors (70-80%) are introduced during sequencing, while the different DNA extraction steps account for at most 30% of these genotypes. In none of our simulations did a genotype with errors occur more than three times (see Figure 3D). Because our simulation showed that a genotype that occurs more than three times in the sequence data is unlikely to be the result of methodological error, we constrained our subsequent analysis to genotypes that are above this threshold. Of the 31’229 distinct genotypes in the sequencing data only 884 genotypes occur more than three times, representing 80’886 individual genomes. In all samples pooled over every evolution line at both time-points and the ancestor population, there are 184 sites in the φX174 genomes that are mutated compared to the consensus. In the ancestral population, there are 53 genotypes and 38 mutated sites. All these statistics are calculated in the file “R-code/code/stats-for-paper-rrr.html” in the supplementary zip-archive “data+codefor-reviewers-Feb25-2026.zip”. A list of all 184 mutations and their characteristics can be found in the supplementary file “mutations.xlsx”.

### Whole-population, whole-genome data show the evolution of mostly new genotypes in each evolution line

The sequencing data that we generated in our evolution experiments is so detailed — especially due to the fact that we have thousands of whole-genome sequences at each sampled time-point — that it presents a challenge for visualization. To overcome this challenge, we present several, partly newly-developed ways to visualize our whole-genome, whole population sequencing data.

A conventional summary plot of the sequencing data for evolution line 1 at generation 412 is shown in Figure 2. This plot shows the frequency of non-consensus nucleotides at each position in the genome. In the sample shown, polymorphisms more frequent than 1:1000 occur in 260 sites across the genome, of which the most frequent occur in the F gene that encodes for the major capsid protein (Figure 2A). This visualization of the sequencing data, however, ignores the linkage of mutations between different sites, and does not show the frequencies of each genotype.

**Figure 2:**
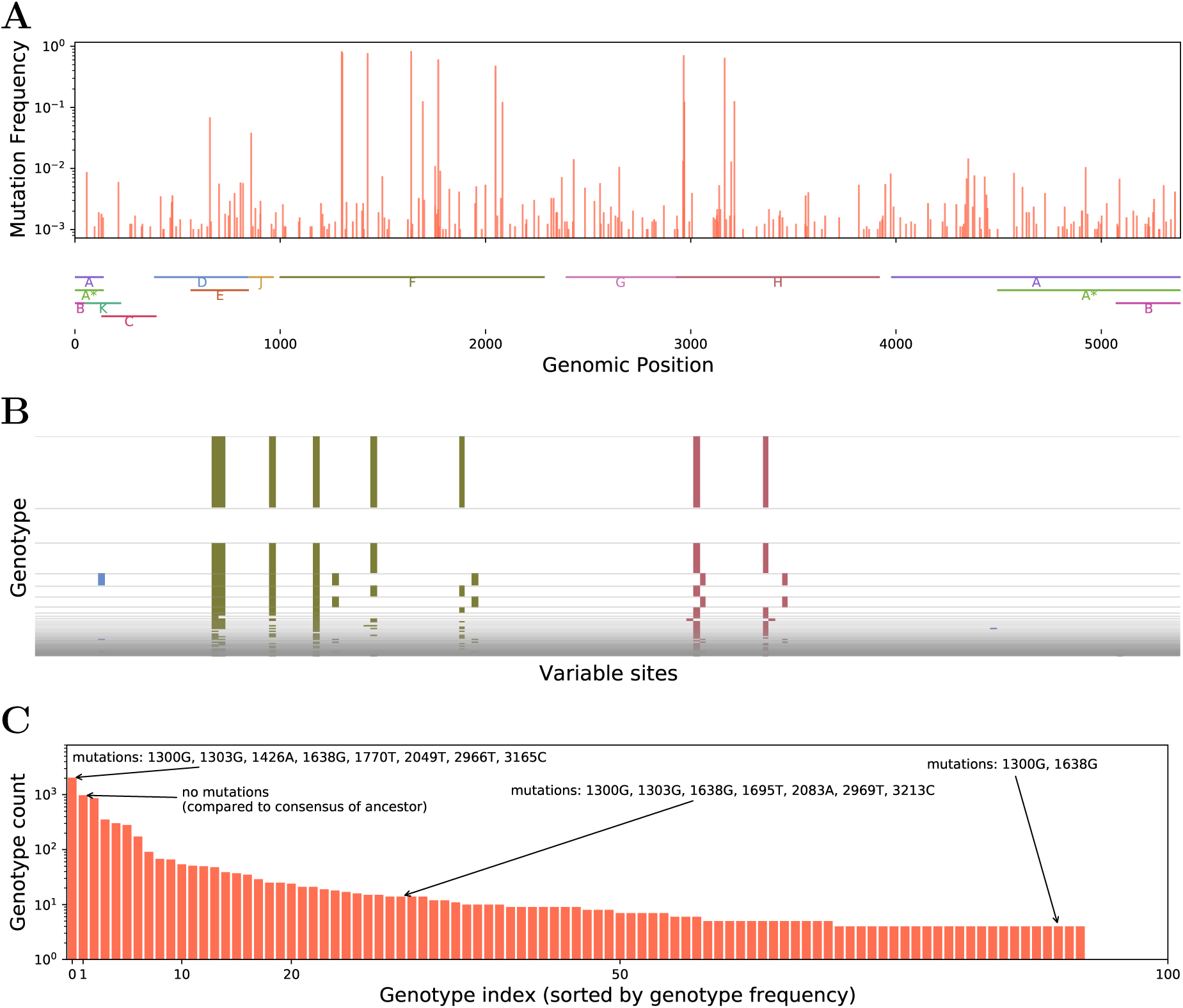
Overview of the viral sequencing data of the first evolution line after 412 generations at different levels of abstraction. **(A)** Frequency of mutations that occurred during the evolution experiment and their location in the genome. The locations of the genes A, A^∗^, B, C, D, E, F, G, H, J, and K of φX174 are indicated below the plot. This representation of the data ignores the genetic linkage between mutations. **(B)** Representation of genotypes present in the population. Each row corresponds to a single genotype, the height of the rows indicate the frequency of the genotypes, The top row displays the most frequent, the second row from the top the second most frequent gentoypes, etc. The x-axis indicates the mutated positions. For the sake of visual clarity, we do show only genomic positions that were mutated at some point in the evolution experiment across all evolution lines and sampling times. Colored vertical lines indicate the positions where the genotype in question was mutated, the color indicates the gene in which this mutation occurs (same colors as in **(A)**). When a mutation occurs at a position with overlapping genes, only one gene is indicated.) This stack of genotypes is the most comprehensive representation, but makes very rare genotypes invisible. **(C)** Genotype count distribution. Genotypes in this plot are indexed by descending frequency in the population (not genetic similarity). This nicely summarizes the number and frequency of distinct genotypes in a sample, but ignores the mutations a genotype carries. (We have annotated some of the genotypes with the mutations they carry to make this clear.) Rare genotypes become visible by using the log-scale, which would be very challenging to implement in **(B)**.

**Figure 3:**
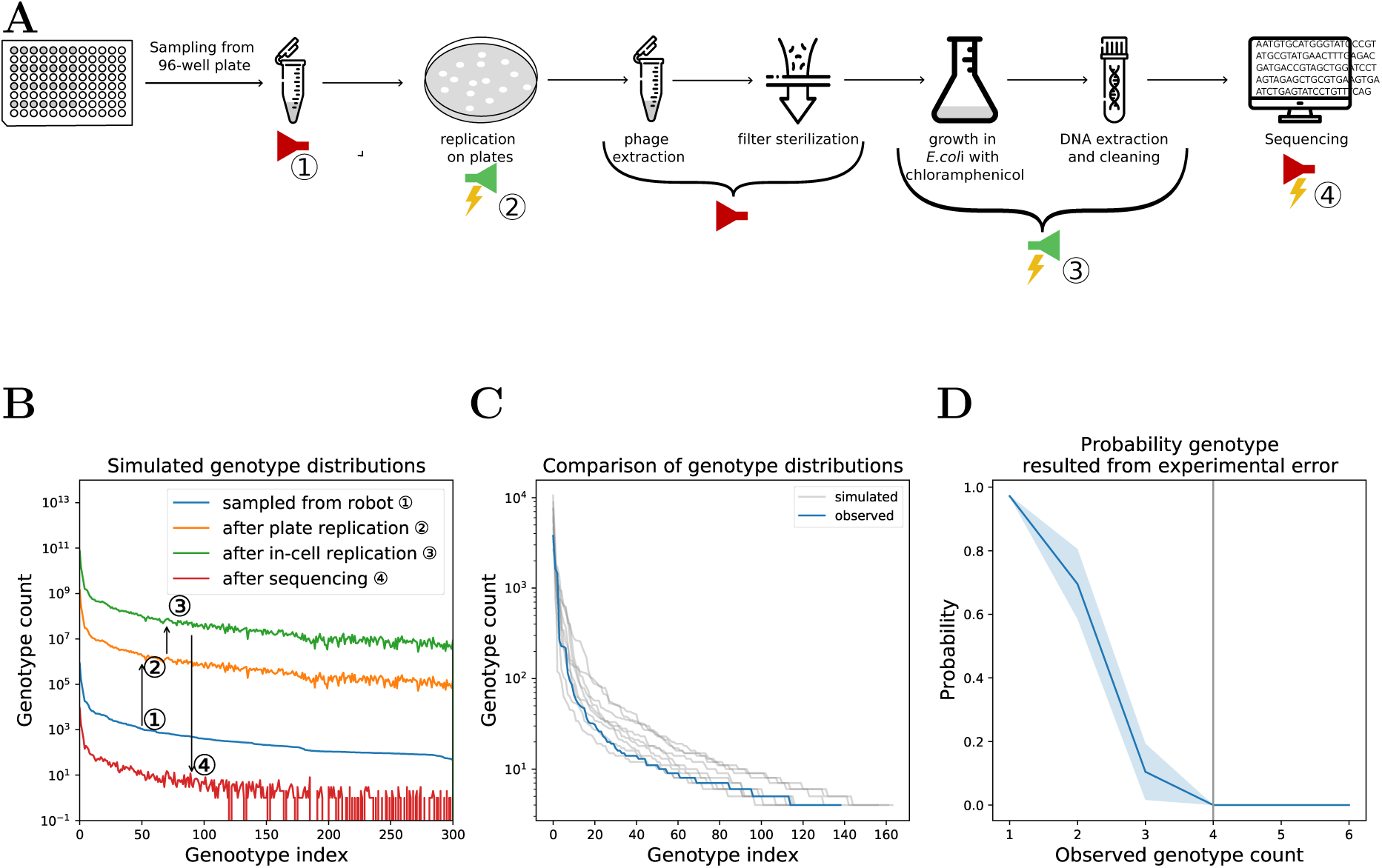
*In silico* simulations of the passaging experiment, DNA extraction, and sequencing. **(A)** Overview of experimental steps involved in the sampling, DNA extraction and sequencing of the bacteriophage population. The red symbols indicate experimental steps that were associated with a population bottleneck (e.g. sampling or filtering) The green symbols indicate experimental steps that were associated with an amplification of bacteriophage populations. Lastly, the yellow lightning bolts indicate experimental step during which mutations could have been introduced. The bottlenecking, amplification and mutation processes were implemented in a detailed simulation model (see Materials and Methods). **(B)** Effect of the different steps of the extraction process on genotype diversity. The circled numbers correspond to experimental steps shown in **(A)**. The first steps increase the population size without significantly affecting the genotype distribution. The final sequencing step forms a major bottleneck, reducing the population size with a factor 10^8^ and causing some genotypes to get lost. **(C)** Comparison of the observed to 10 simulated genotype distributions. The simulated genotype count distributions are very consistent with the observed one. **(D)** Probability that a genotype observed after sequencing (step ○**4**) was present in the original population at step ○**1**,based on 10 independent simulations of the experiment and DNA extraction. Genotypes that are present more than 3 times in the sequenced population were very likely present in the original population.

Figure 2B displays the information on the diversity and linkage in a bacteriophage population sample by showing stacks of genotypes. This is similar to highlighter plots as used in the context of HIV, e.g. by Keele et al. (2008). The height of these stacks is proportional to the frequency of the genotype in the sample. The genotypes are displayed as a reduced genome, that contains only the sites that are polymorphic. (In this regard this plot differs from the highlighter plot that typically displays the entire genome.) This visualization, while containing information on the frequency of each genotype, does not reveal low frequency genotypes. The sample of the virus population in evolution line 1 at generation 412 that we display in Figure 2B consists of 6187 genomes, representing 93 distinct genotypes. The most prevalent genotype was observed 2035 times, the second most prevalent — the consensus in the ancestor population — was found 973 times. 23 genotypes in this sample were found 4 times (the lowest frequency above the cut-off defined above). These low frequency genotypes are not visible in Figure 2B.

To be able to visualize the full frequency distribution of genotypes in a sample, we developed what we refer to as *genotype count distribution* (Figure 2C). In this plot distinct genotypes are indexed in order of decreasing frequency, and the logarithm of their frequency is shown as a histogram. This clearly displays the number and frequency of each genotype in a sample, but, unlike a highlighter plot, does not display the specific mutations each genotype carries.

Using these genotype count distributions, the diversification of the bacteriophage population over time can be visualized in great detail (Figure 4). The grey genotype count distributions at the left of each evolution line in Figure 4 show the ancenstor population, which is used to seed every evolution line and consist of 53 distinct genotypes. The lightly colored genotype count distributions in each line show the composition of the bacteriophage populations at generation 196. They largely do not overlap with the ancestor population, indicating substantial shifts in the viral composition over time in each evolution line. There is also almost no overlap between the genotype count distributions between the evolution lines. Thus, most genotypes that evolve are new and unique to the evolution line in which they evolved — as would be expected in a regime of neutral evolution.

**Figure 4:**
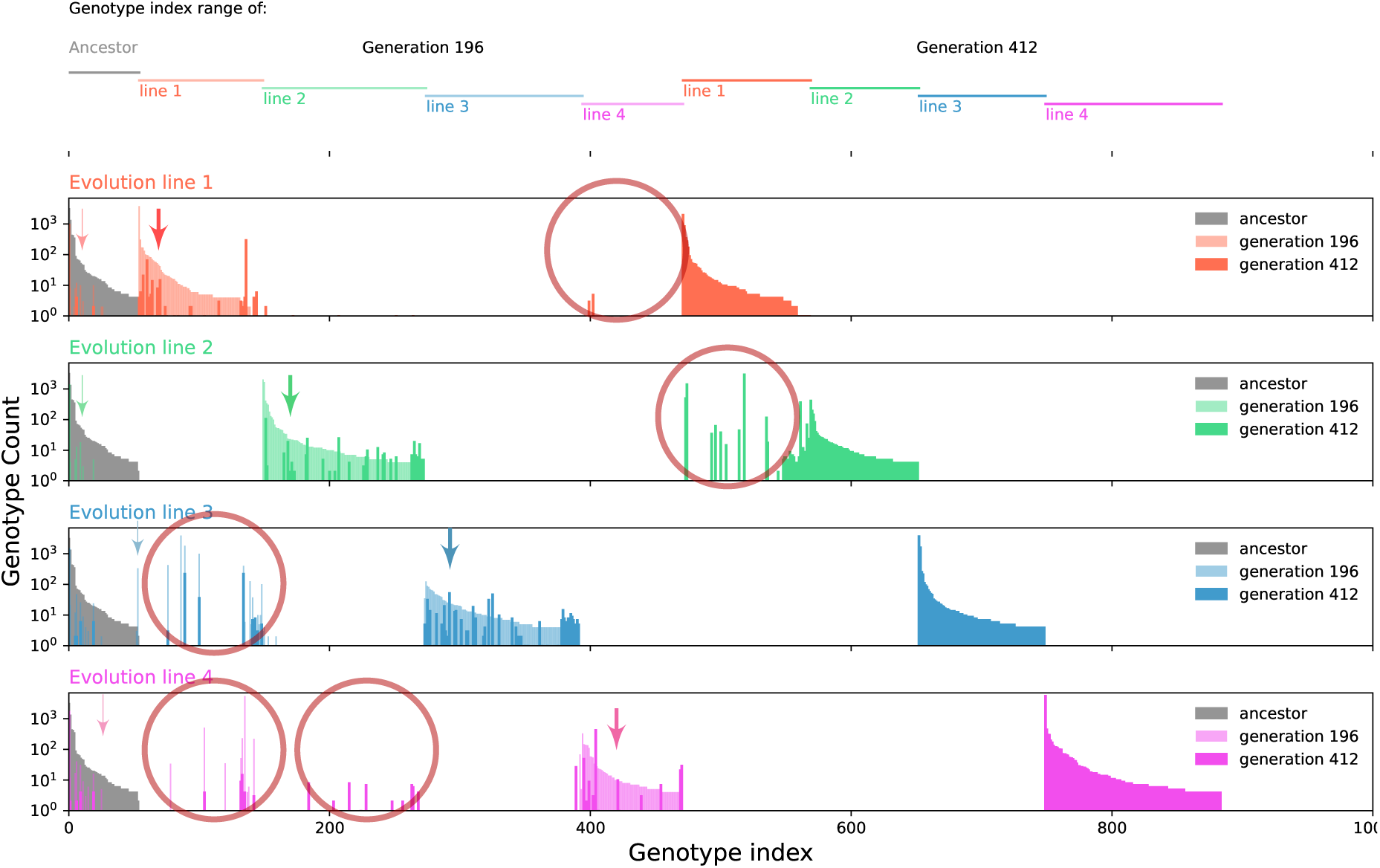
Diversification of the φX174 populations in each evolution line over time. Each row represents an evolution line, and shows the genotype count distributions for the ancestral (grey) and evolved viral population after 196 (light colors) and 412 (dark colors) generations. As in Figure 2C, we indexed the genotypes within the viral population at a given time-point by descending order of frequency in the sample. The grey distributions on the left of each line show the counts of the 53 distinct genotypes in the ancestral population. The ancestral genotype count distribution is the same in each evolution line. New indices were added for genotypes in subsequent samples that did not occur, and were thus not indexed, in a previous sample. An overview of the index ranges for each evolution line and time-point is shown at the top of the graph. The shift of the genotype count distributions to the right in each line means that most of the genotypes differ between the viral populations from different time points. The low degree of overlap in the genotype count distributions across evolution lines is due to the fact that the newly evolved genotypes are largely unique to an evolution line. Not all genotypes are unique to the line and time-point, however. Arrows point to examples of genotypes from the ancestral and mid-point populations that are still present in later samples. The red circles highlight genotypes that evolved in parallel in multiple evolution lines. Note these instances of evolution are parallel on the level of the whole-genome, not just on the level of mutations at single sites in the genomes.

An alternative visualization of the evolution of the phage in the four lines is the genotype network shown in Figure S3. While this graph shows only a subset of the 50 most prevalent genotypes, it clearly illustrates the independent evolutionary trajectories in each line.

### Extensive parallel evolution of single mutations, combinations of mutations, and even whole-genome genotypes

Although, we generate mostly unique genotypes over time in each evolution line in our passage experiment, there are genotypes that emerged in multiple evolution lines (indicated by red circles in Figure 4). These genotypes are very unlikely to have arisen by cross-contamination during passaging the bacteriophage. This is because, first, on the 96-well plate we used for passaging, we separated the wells in which the bacteriophage was passaged by control wells that were seeded with bacteria to detect spill-over (see Materials and Methods). Most of these control wells were not infected by spilling-over bacteriophage, and in case of infection, the experiment was restarted (see Materials and Methods), making it unlikely that spill-over occured across two rows on the 96-well plates. Second, many of the genotypes that emerged in multiple lines were even further apart on the 96-well plate (quite a few between evolution lines 1 and 3, as well as 1 and 4 - see Figure 4). Lastly, the parallel genotypes are never the most frequent genotypes in any of the evolution lines in which they appeared (Figure 4) Thus these genotypes most likely represent instances of parallel evolution.

It is important to emphasize that these instances of parallel evolution represent the parallel evolution on the level of whole genomes, rather than parallel evolution of single mutations. In total, we observe 26 genotypes that evolve in more than one evolution line. Compared with the ancestor genotype that is genetically closest, they aquired 1 to 5 additional mutations (mean 2.5). In total, these 26 genotypes carry 18 mutations that are not found in the ancestor population (see the file “R-code/code/stats-for-paper-rrr.html” in the supplementary zip-archive “data+code-for-reviewers-Feb25-2026.zip”). Of these mutations 33% are synomymous, compared to 19% in the ancestor, and 25% in the entire dataset. Establishing parallelism on the whole-genome level requires the whole-population, whole-genome sequencing approach presented here that also captures genotypes of low frequency.

To investigate to what extent the observed parallel evolution is consistent or inconsistent with an accumulation of unbiased and selectively neutral mutations, we performed randomization tests (see Materials and Methods). Specifically, we compared the frequency of the observed parallel individual mutations and combinations of mutations with the frequency predicted under unbiased neutral evolution. (Figure 5).

**Figure 5:**
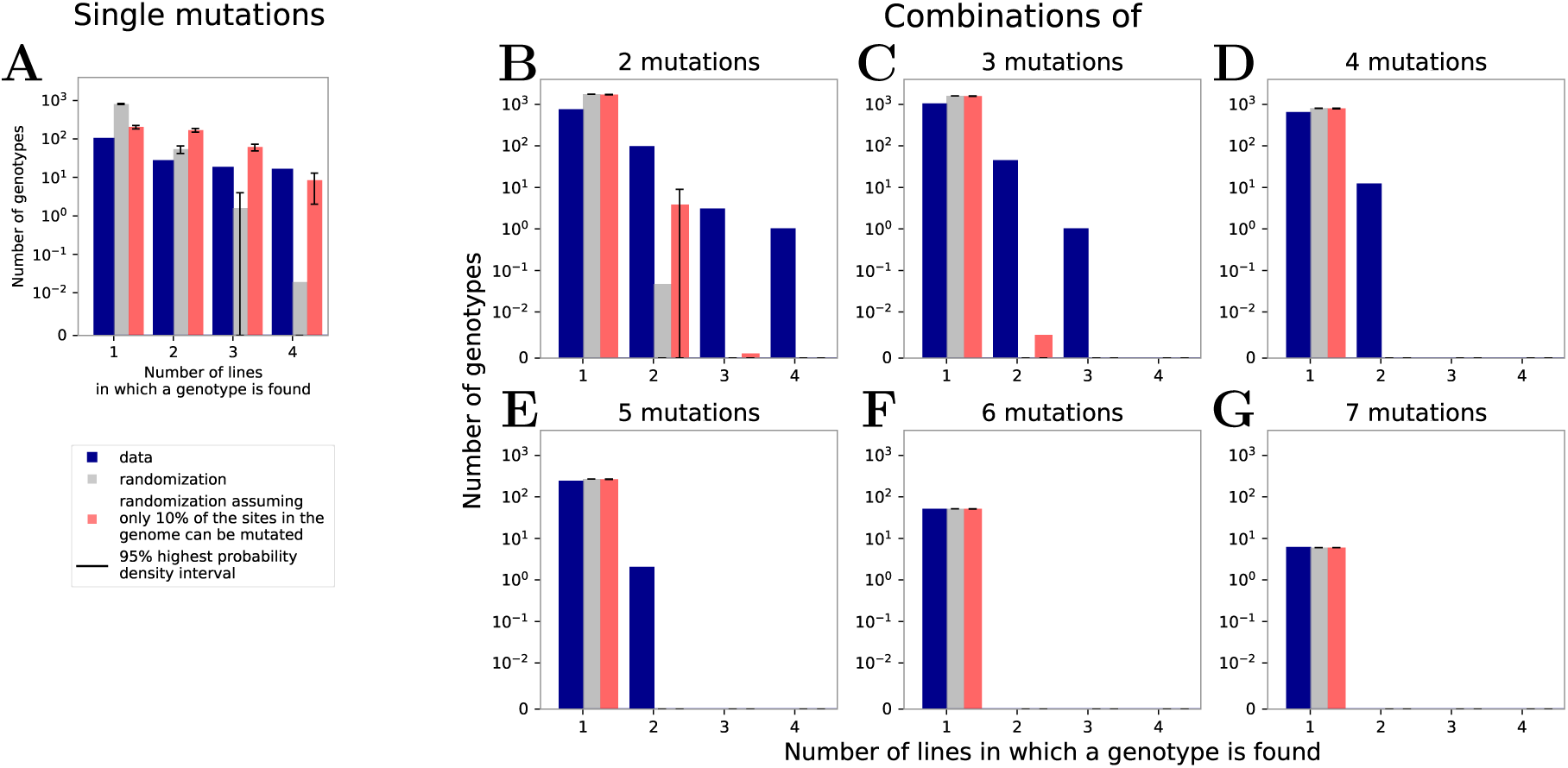
Parallel evolution exceeds the level predicted by a model of neutral evolution. The plots show the number of single mutations **(A)**, and combinations of mutations **(B–G)** against the number of evolution lines in which they appear. A mutation that occurs in more than one evolution line represents an instance of parallel evolution. The blue bars show the frequencies of mutations or their combinations observed in the experiments. The grey and red bars show expectations based on 10’000 randomizations (see Materials and Methods). We observe the parallel evolution of combinations of up to five mutations. The randomizations indicate that under neutral evolution combinations of more than two mutations are unlikely to emerge in more than a single evolution line.

On the level of individual mutations, we found 137 that emerged in only a single evolution line, while 47 emerged in multiple lines. The fraction of synonymous mutations does not differ significantly between parallel and non-parallel mutations (see the file “R-code/code/stats-for-paper-rrr.html” in the supplementary zip-archive “data+code-for-reviewers-Feb25-2026.zip”). Seven mutations were shared in all four evolution lines (Figure 5A - blue bars). Randomization tests show that this level of parallel evolution is not consistent with unbiased, selectively neutral mutations (Figure 5A - gray bars). This extent of parallelism is similarly high as in previous evolution experiments using the same bacteriophage (Bull et al., 1997, Wichman et al., 1999, 2005) or other viruses (Bertels et al., 2019, Bons et al., 2020).

The linkage information in our data puts us into the unique position to investigate the parallelism of combinations of mutations across the entire genome (Figure 5B–G). We observed the emergence of combinations of up to five mutations in multiple evolution lines (Figure 5E). Randomization tests show that the extent of parallelism is significantly higher than that predicted by neutral evolution. Randomization tests with an effectively shorter genome (see Materials and Methods) show that the parallelism of combinations of mutations cannot be fully explained by the excess parallelism of single mutations. This strongly suggests that the selective advantage of the mutations in combination must on average exceed that of the individual mutations involved, i.e. the mutations in many of the parallel-evolved combinations are positively epistatic. More direct evidence for positive epistasis between mutations could be gained from more fine-grained data on the frequency of genotypes over time. This, however, will require to sample and sequence more frequently than every 200 generations.

Whether these combinations arose by successive point mutations, or by recombination is difficult to ascertain. The multiplicity of infection during the passaging experiment was on the order of 1, making it likely that bacterial cells were co-infected, enabling recombination. An analysis of genetic linkage (see Materials and Methods), however, did not find any evidence for signatures of recombination.

### Phylodynamics inference methods over-estimate migration rates between evolution lines

Viral sequences are widely used for phylodynamics inference which in turn has become key to study the spread and evolution of important viral pathogens between-host (Fraser et al., 2009, Faria et al., 2014, Gire et al., 2014, Düx et al., 2020, Nadeau et al., 2021), as well as within-host (Lorenzo-Redondo et al., 2016). However, parallel evolution can lead to biases in phylodynamic inference (Koelle and Rasmussen, 2025, Ochsner et al., 2026).

To investigate to what extent the parallel evolution we observed in our experiment biases parameters estimates, we conducted a phylodynamic analysis, estimating the migration rates between the independent evolution lines (see Materials and Methods). Because the viral populations evolved independently in each line, there is no migration between the lines, and the estimated migration rates should not be significantly different from 0.

Figure 6A shows the distribution of estimated overall migration rates between evolution lines. In this distribution, the migration from the ancestor to the evolution lines has been subtracted. In a nutshell, the migration rate estimates are significantly different from zero, which they should not be because there is no migration between the evolution lines in the experiment. The point estimate is 0.00275 per generation, which corresponds to a per generation probability of 0.275% for a genome to migrate from one evolution line to the other. Over the course of the 412 generation experiment, this migration rate amounts to a 68% probability for each single genome to migrate to a different evolution line. Qualitatively, these insights also hold when we exclude the 26 genotypes that evolved in parallel Figure 6B.

**Figure 6:**
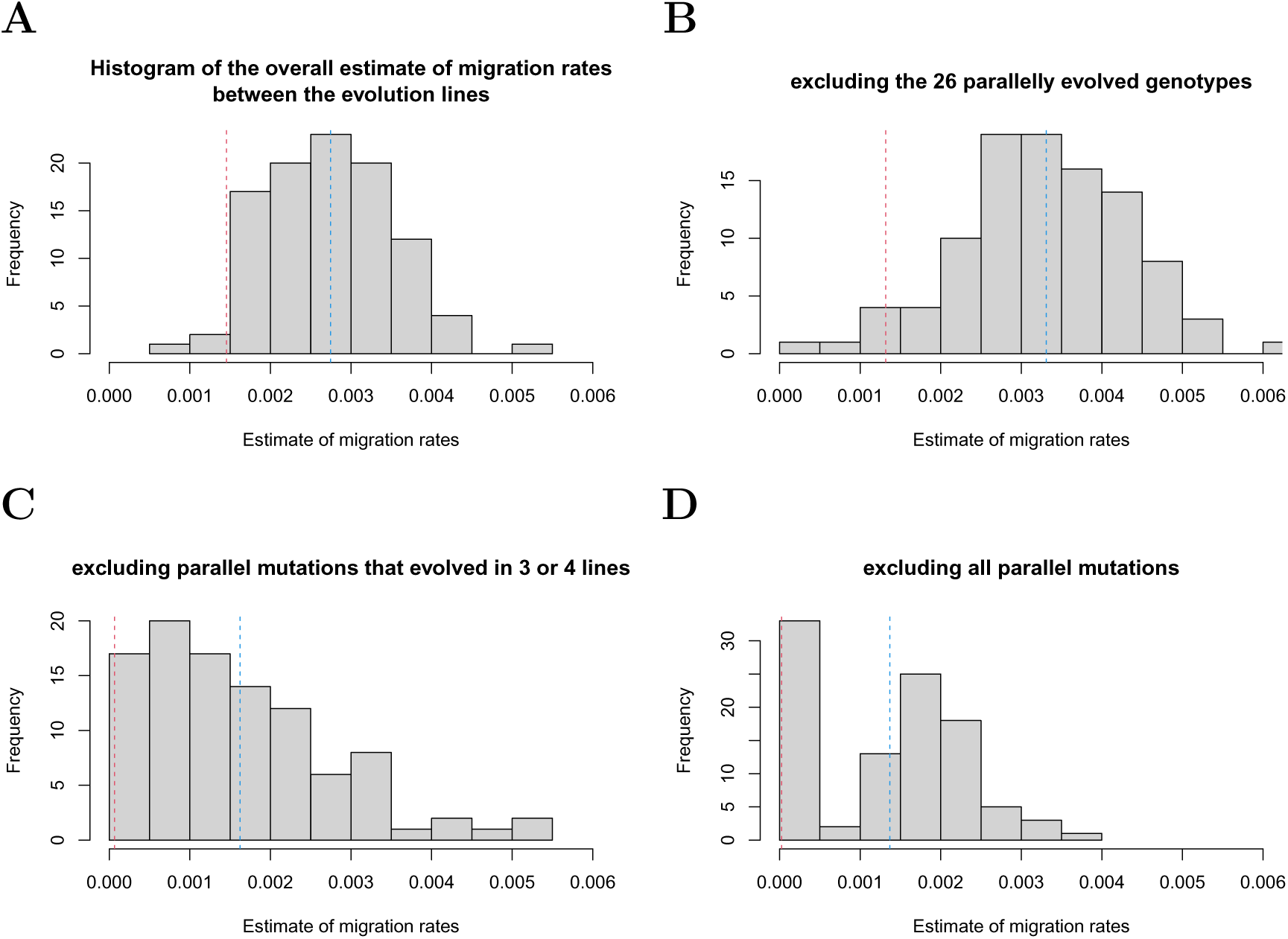
Distribution of migration rate estimates from the treetime analysis. The red dashed line shows the lower bound of the 95% confidence interval of the estimates, and the blue dashed line their mean. **(A)** The estimates were generated by subsampling 900 sequences from the total of 80’886 whole-genome sequences we obtained in the experiment, reconstructing the phylogeny, and performing the mugration analysis (Lemey et al., 2009) implemented in treetime (Sagulenko et al., 2018). Migration rates between the ancestor population and the evolution lines have been subtracted from the overall migration rate estimate to obtain an estimate of the overall migration between the evolution lines only (see Materials and Methods). **(B)** Same analysis as in (A) except that we excluded from the data set the 26 genotypes that evolved in parallel in multiple line. In both cases, (A) and (B), the overall migration rate between the evolution lines is wrongly estimated to be significantly larger than zero. **(C) and (D)** same analysis as (A) and (B), but here we masked excess parallel mutations (C), i. e. mutations that evolved in three or four evolution lines, or all parallel mutations (D). Masking parallel mutations corrects the bias in migration rates.

This positive migration rate, rather than reflecting real migration, constitutes a bias that arises because the analysis does not account for the selection pressures that generate the patterns of parallel evolution in our experiment. Thus, even in systems as ours, in which no intentional selection pressure is imposed on the viral population, selection is strong enough to bias phylodynamic inference.

To provide direct evidence for the role of parallel evolution in the over-estimation of migration rate, we conducted a phylodynamic analysis masking all mutations that appeared in multiple lines, i.e. parallel mutations, or just those that appeared in 3 or 4 lines which we call “excess parallel mutations” because only these are more numerous than neutral simulations suggest (see Figure 5A). We find that, in both case, the lower bound of the 95% confidence interval of the migration rate estimate is not above 0 anymore, meaning that the bias is not statistically significant anymore. This clearly shows that parallel mutations are the source of the overestimation bias.

## Discussion

In summary, we present an approach to capturing the molecular evolution of viruses with short genomes in unusually high detail. The approach gives rise to very accurate (error rate per nucleotide *<* 2×10^−5^) sequence reads spanning the entire genome without the typical loss of linkage information associated with short-read sequencing methodology. With approximately 9000 whole-genome reads per sample on average, we obtain fine-grained information on the frequency of each genotype throughout the 400 generations evolution experiment. To our knowledge, sequence data at this level of detail and accuracy do not exist for any organism to-date.

The picture of bacteriophage evolution we document is characterized the emergence of mostly new genotypes in each line that are unique to that line. But we also established instances of significant parallel evolution of many mutations and their combinations, and even of 26 genotypes that were identical across the entire genome.

We found that phylodynamic analysis wrongly infers significant migration between the independent evolution lines. We chose to conduct these analyses with treetime because it handled our big dataset with its many polytomies well. But the biases we identify should not be attributed to problems with treetime’s estimation of migration rates. They arise in phylodynamic inference schemes in general due to parallel evolution (see the discussion below).

The inaccuracy of the reconstructed phylogenetic tree could be quantified by the number of times a lineage assignment switches from one evolution line to another. This quantity, however, is not very different from the migration rate estimate which, in a nutshell, is derived from the number of such switches divided by the tree height (Lemey et al., 2009). Thus, the distributions of migration rate estimates in Figure 6 can also be interpreted as the distribution of the inaccuracy of the reconstructed phylogenetic trees. As direct evidence that parallel evolution leads to the biases in phylodynamic inference, we conducted analyses with datasets that either did not include the 26 genotypes that evolved in multiple lines, or masked (excess) parallel mutations. While the analysis without the 26 parallel genotypes did not correct the bias, masking parallel mutations did (see Figure 6). Although this analysis clearly shows that parallel mutations are the cause of the bias, it is not a viable solution to the problem because parallel mutations are difficult to identify. In our experiments we did not impose any selection intentionally, and most of the parallel mutations were unanticipated and unknown. In epidemiological systems with potential migration between locations it will be extremely challenging to disentangle whether mutations co-occur in multiple locations due to parallel evolution or migration. Exceptions are known antiviral drug-resistance mutations that are commonly masked when reconstructing the transmission history of HIV-1.

Phylodynamic inference methods are usually tested on simulated data. The simulation models often conform with the assumptions of the inference method, most importantly that of neutral evolution (for note-worthy exception see (Bedford et al., 2011, Rasmussen and Stadler, 2019)). There are only few studies that generated data that were biologically more realistic (Ratmann et al., 2017, Ochsner et al., 2026). A comprehensive assessment of phylodynamic methods sometimes would require testing their performance on simulated data that, rather than conform with the assumption of phylodynamics, capture important biological, demographic, and epidemiological aspects of the system in question. The reason we resorted to an experimental system, rather than simulations here, is because non-neutral evolution is challenging to simulate. We simply know too little about how viral fitness changes with each mutation, how effects of each mutation combine, and how the composition of the viral population affects selection. The surprisingly high degree of parallel evolution we observed in this study — despite not intentionally imposing any selection pressure on the phage — is a case in point.

There is precedent to experimentally testing population genetic methods that reconstruct evolutionary history from genetic data. Experimental evolution with phages has been employed to test phylogenetic methods for over three decades (Hillis et al., 1992, Bull et al., 1997, Turner et al., 2012). The results were mixed. Hillis et al. (1992) used the phage T7 and generated restriction-site maps, not genetic sequences, and could infer phylogenetic trees with the correct topology but not branch length, while Bull et al. (1997), using the phage φX174 that we also used in the present study, found that phylogenetic reconstruction failed because of “exceptional convergent evolution”. In the more recent study by Turner et al. (2012) with the phage φ6, the evolutionary history could not be recovered by some phylogenetic methods because of a lack of viral diversity accumulation in their evolution experiments. We see our work in this tradition, extending it to the recent developments in phylodynamics that use the inferred evolutionary histories to estimate population biological parameters.

Similar to the landmark paper on the experimental evolution of φX174 (Wichman et al., 1999), we could not prevent recombination. The multiplicity of infection in our experiments was at 0.08 at the beginning of the evolution experiment and increased to 2.8 by its end, suggesting that co-infection of bacterial cells by phages — a prerequisite to recombination — was increasingly likely. Thus, we cannot exclude that recombination occurred. Recombination, however, does not always give rise to recombinant variants, especially if the sequence diversity of the evolving population is low. This is the likely reason for the finding that an linkage disequilibrium analysis did not find a significant recombination rate.

Recombination poses a problem for any analysis that relies on reconstructing the evolutionary history, such as phylodynamics. In epidemiological settings recombination often occurs (Holmes, 2009, Müller et al., 2022), and the incorporation recombination has been recognized as one of the major challenges in phylodynamic inference (Frost et al., 2015). In our experimental setting, however, recombination is unlikely to significantly distort the inferred evolutionary histories because the genotypes in our experiment are highly related, and correct phylogenies can be successfully inferred in such a regime (Posada and Crandall, 2002). This suggests that the biases we identify are not caused by recombination per se, but by the parallel selection pressures between the evolution lines.

The fact that we cannot rule out recombination in our experiment is less of an issue for applications of phylodynamic methods to the viral population dynamics within infected hosts, referred to as *phyloanatomy* (Salemi and Rife, 2016, Lorenzo-Redondo et al., 2016). As recombination rates of viruses within-host can be high (Neher and Leitner, 2010, Schlub et al., 2014, Romero and Feder, 2024), our experimental system is an appropriate reflection of the within-host dynamics of viruses. Even if recombination distorts the inferred within-host phylogenies (which we do not believe is the case in our experiments — see above), the resulting potential biases of phylodynamically-inferred migration rates and other population parameters would very likely reflect biases we could expect in the within-host context.

In addition to the detailed insights into the long-term viral evolution and the validity of phylodynamics inference, we believe that the approach we present here will open new avenues to trace viral population genetics on the whole-genome level, i. e. observe the emergence, drift and selection of mutations in combination with the genetic background on which they appear. With sufficiently frequent sampling and sequencing, it will also allow the tracking of the temporal dynamics of the many concurrent genotypes in virus populations, and the direct observation of complex phenomena, such as clonal interference. Such temporally highly-resolved, whole-genome, whole-population data will enable better estimation of the fitness of the viral genotypes, as well as main and epistatic effects of mutations that are at the heart of population genetics.

## Materials and Methods

### Strains, growth conditions and bacteriophage titration

Except otherwise stated, *Escherichia coli* C (ATCC 13706) is cultured in LB Miller broth (Sigma-Aldrich, Cat. L3022) at 37^◦^C with aeration at 180 rpm. For infection with bacteriophage φX174 (ATCC 13706-B1, obtained from the DSMZ collection), the medium was supplemented with 10 mM MgCl_2_ and 2 mM CaCl_2_.

Amplification of φX174 was carried out by inoculating 50 mL of supplemented LB broth with 500 *µ*L of stationary-phase *E. coli* and 250 *µ*L of φX174 stock. Cultures were incubated overnight at 37^◦^C with aeration. Following incubation, the cultures were centrifuged to pellet bacterial cells, and the supernatant was filtered through a 0.22 *µ*m filter to isolate the phage-containing lysate.

Bacteriophage titration on plates was carried out by plaque assay. Specifically, 6 mL of soft agar, LB Miller soft agar (LB broth containing 5 g/L agar (Sigma-Aldrich, Cat. A7002), 10 mM MgCl_2_ and 2 mM CaCl_2_) was mixed with 400 *µ*L of stationary-phase bacteria and 20 *µ*L of diluted φX174 lysate, then poured onto LB Miller hard agar plates from LB broth with agar (Sigma-Aldrich, Cat. L2897) and the same salt concentrations. Plates were incubated overnight at 37^◦^C in an inverted position, and plaques were counted to determine the initial phage concentration.

For liquid-based titration, a 96-well plate format was used. Each well was filled with 200 *µ*L of LB Miller broth containing 1% (v/v) stationary-phase *E. coli* and φX174 diluted to an average of 0.5 plaque forming units (PFU) per well. Plates were incubated overnight at 37^◦^C, and the presence of phage replication was assessed by spotting 2 *µ*L from each well onto a fresh bacterial lawn prepared identically to those used in the plaque assay but without added phage. After a 5-hour incubation at 37^◦^C, wells with no evidence of phage replication were counted. The initial phage concentration was estimated by calculating the fraction of wells without detectable phage replication, assuming a Poisson distribution of phage particles across wells.

### Experimental evolution of bacteriophage φX174

Bacteriophage phiX174 was experimentally evolved for 412 generations in 4 independent evolution lines. The experimental evolution of bacteriophage φX174 was carried out in 96-well plates using a Tecan Freedom Evo 200 automated liquid handling system (Tecan) with an integrated automated incubator (Liconic STX100, Liconic). At each transfer, 4 *µl* of supernatant was passaged into the next well.

The passaging terminated in the liquid handling system 3 times at generation 95, 150 and 309. At generation 95, only 2 *µl* of supernatant was passaged.

Bacteriophages were passaged from column to column on a 96 well-plate approximately every 26 minutes, allowing for one complete replication cycle in each well (Baker et al., 2016). On a single column, 4 replicated lines of bacteriophage φX174 evolve, each separated by a line containing only host bacteria. This allows the detection of contamination events: a delay in bacterial growth in these lines indicates a contamination by the bacteriophage. For each generation n, we realise a transfer (described in the next paragraph) and then incubate the plates for 20 minutes at 37^◦^C and 95% relative humidity. Once a day, we make a glycerol stock by mixing 100 *µ*L of 40% glycerol with 100 *µ*L of the n-1 bacteriophage generation. The stocks are then stored at -80^◦^C. In case of bacteriophage contamination or other problems, the experiment is restarted using the last stock as a bacteriophage source.

Each transfer is composed of several steps with pipette sterilisation after each step (see below). First, on column n+6, 5 *µ*L of freshly grown overnight *E. coli* C are used to inoculate 200 *µ*L of LB Miller supplemented with 10 mM MgCl_2_ and 2 mM CaCl_2_. Second, 4 *µ*L of freshly mixed column n-1 is transferred into column n. As a result, bacteria are inoculated approximately 1h30 before bacteriophage infection: this assures that bacteriophage infection occurs on exponentially growing bacteria. In addition, this protocol aims to ensure that the multiplicity of infection stays largely below one, which limits bacteriophage recombination. Third, if needed, glycerol stock of bacteriophages are made as previously described.

The entire experiment is carried out with reusable tips, which require sterilisation between each step to inactivate any bacteria or bacteriophage particle. To do so, the tips are washed twice in 5.6% bleach, flushed from behind with water, washed in 70% ethanol and then in milliQ water.

To calculate the multiplicity of infection (MOI), we estimated the density of *E. coli* and φX174 in a well. We calculated *E. coli* density by performing a standard colony forming unit calculation as follows: 5 *µ*L of stationary phase bacteria are inoculated into a well with 200*µ*L of LB Miller. After 90 minutes of incubation at 37^◦^C, a serial dilution is prepared from the culture to back-calculate the density in the well. 100*µ*L of each dilution are then spread onto a petri dish containing Hard Agar (LB Miller with 15 g/L of Agar, 10 mM MgCl_2_ and 2 mM CaCl_2_). The plate is incubated upsidedown overnight at 37^◦^C. We counted bacterial colonies and determined that there were 1.84 × 10^7^ bacteria/well. We calculated φX174 abundance using the plaque-forming unit protocol described above at generations 0, 196 (midpoint), and 412 (endpoint). Here, we report measured PFU of phage in 4*µ*L as this is the volume used to initiate the experiment and transfer between wells each generation. These were: 1.44 × 10^6^ PFU at generation 0, 2.42 × 10^7^ PFU at generation 196 and 5.12 × 10^7^ PFU at generation 412. Combining these PFU measurements with the number of bacteria per well we obtain MOIs of 0.08, 1.38, and 2.8 at generation 0, 196, and 412, respectively.

### Extraction of the φX174 DNA

Two protocols were developed to extract φX174 DNA from experimental samples, targeting different stages of the bacteriophage’s life cycle (dsDNA vs. ssDNA) and using distinct amplification and extraction strategies. (New protocols were needed because PacBio long-read sequencing requires 1 *µ*g of phage DNA, many orders of magnitude more than we have in each well during our experiment.) The dsDNA extraction protocol was applied to all samples, forming the basis of the main dataset.

To compare the protocols, both methods were applied to two samples. As shown in Figure S2, the dsDNA protocol yielded a greater diversity and abundance of distinct genotypes. Due to its higher efficiency, it was selected as the standard extraction protocol for the study.

#### dsDNA extraction

A ready-to-use version of this protocol is available in the supplementary information.

φX174 DNA was extracted during the replication stage when the genome forms a double-stranded circular dsDNA intermediate. This replication intermediate was stabilized by chloramphenicol-induced inhibition of bacterial protein synthesis cycle (Sinsheimer et al., 1962, Denhardt and Silver, 1966).

First, high-titer lysates were generated by amplifying 10% of the phage population in each sampled well on agar plates. Plates were overlaid with soft agar containing 400 *µ*L of stationary-phase *E. coli* and 20 *µ*L of phage sample. After overnight incubation at 37^◦^C, phages were extracted by adding 3 mL of LB broth to the plate, gently mixing, and recovering 2 mL of the supernatant, which was then filtered through a 0.22 *µ*m filter.

Second, 200 ml cultures of *E. coli*, infected with φX174 lysates, were incubated with chloramphenicol and harvested after 3 hours. To prevent recombination during phage replication, the multiplicity of infection was kept below 1 (see validation below).

Third, DNA was extracted using a modified protocol based on the plasmid miniprep protocol (NEB, Cat. T1010), adapted for larger culture volumes and sequential spincolumn loading to isolate phage-derived double-stranded DNA.

Last, the extracted DNA was then linearized using NEB Cutsmart Buffer (NEB; Cat. B7204) and PstI-HF restriction enzyme (NEB, Cat. R3140) and finally purified with AMPure XP (Beckman Coulter, Cat. A63880).

DNA quality and quantity were assessed using three methods: concentration was measured with the Qubit dsDNA BR Quantification Assay (Thermo Fisher Scientific, Cat. Q32850) on a Qubit Flex Fluorometer, chemical purity was evaluated with a Nanodrop spectrophotometer (Thermo Fisher Scientific), and fragment integrity and contamination were checked using an Agilent TapeStation with Genomic DNA ScreenTape (Agilent Technologies, Cat. 5067), following the respective manufacturer instructions.

To validate that the MOI remained below 1 during infection, phage and bacterial concentrations were monitored. Phage titers from two samples were determined after overnight amplification on agar plates and found to be 1.2 × 10^9^ PFU/mL and 4.4 × 10^8^ PFU/mL, respectively. Bacterial concentration after 1.5 hours of growth in the 200 mL cultures was measured at 2.94 × 10^7^ CFU/mL, yielding a total of 5.88 × 10^9^ CFU per flask. Given that no more than 2 mL of phage lysate was added per 200 ml flask, the resulting MOI was confirmed to be smaller than 1.

#### ssDNA extraction

A ready-to-use version of this protocol is available in the supplementary information.

To isolate φX174 single-stranded DNA (ssDNA), we used a five-step protocol based on established precipitation, enzymatic digestion, and whole-genome amplification steps.

First, high-titer lysates were generated as described above.

Second, 1 mL of lysate was mixed with 500 *µ*L of 30% PEG/1.5 M NaCl, incubated at 4^◦^C for 24 hours to precipitate the phage particles, and centrifuged twice to pellet the phages. The pellet was resuspended in DNase buffer (NEB, Cat. B0303S), treated with DNase I (NEB, Cat. M0303S) and RNase A (Thermo Fisher Scientific, Cat. EN0531) to remove host nucleic acids and heated to 75^◦^C for 15 minutes to inactivate the DNase. Capsids were lysed by incubating with 20% SDS and pronase (Sigma-Aldrich, Cat. P5147) at 40^◦^C for 2.5 hours.

Third, DNA was purified by ethanol precipitation. 0.5M NaCl and two volumes of cold ethanol were added, followed by overnight incubation at -20^◦^C. The DNA was pelleted, washed with 70% ethanol, dried, and resuspended in 100 *µ*L of DNase-free water.

Fourth, whole-genome amplification was performed using Q5 polymerase (NEB, Cat. M0491) with φX174-specific primers (forward: 5’-ATC GCT TCC ATG ACG CAG AA-3’; reverse: 5’-TGC AGG TTG GAT ACG CCA AT-3’). Each 50 *µ*L PCR reaction contained 21.5 *µ*L nuclease-free water, 10 *µ*L Q5 reaction buffer (NEB, Cat. M0491), 2 *µ*L DMSO, 2.5 *µ*L of each primer (10 *µ*M), 1 *µ*L dNTPs (10 mM), 0.5 *µ*L Q5 polymerase, and 10 *µ*L template DNA. Five parallel reactions were run per sample using the following program: 98^◦^C for 30 seconds; 32 cycles of 98^◦^C for 10 seconds, 68^◦^C for 30 seconds, and 72^◦^C for 108 seconds; followed by a final extension at 72^◦^Cfor 2 minutes. Successful amplification was confirmed using an Agilent TapeStation with Genomic DNA ScreenTape (Agilent Technologies, Cat. 5067).

Finally, PCR products were pooled, purified with 0.8 volume AMPure XP beads (Beckman Coulter, Cat. A63880), according to manufacturer instructions, and eluted in 80 *µ*L of EB buffer (NEB, Cat. T1016) pre-warmed to 50^◦^C. DNA concentration and purity were assessed with a Qubit BR dsDNA assay (Thermo Fisher, Cat. Q32850) and a Nanodrop spectrophotometer.

### PacBio sequencing

The DNA is sent for long-read sequencing to the Functional Genomics Center in Zürich. Sequencing was performed on a PacBio Single molecule sequencer Sequel SMRT Cell 1M.

The long-read sequence data generated in the context of this study have been deposited in the European Nucleotide Archive (ENA) at EMBL-EBI under accession number PRJEB80636 (https://www.ebi.ac.uk/ena/browser/view/PRJEB80636), and will be made publicly available upon acceptance.

The supplementary zip-archive “data+code-for-reviewers-Feb25-2026.zip” contains a file “data/haplotypes run2.csv” that contains the genotypes and their frequencies in all samples extracted from the long-read sequencing data. This dataset retains all the information in the aligned long-read sequences in an efficient way, and is the main dataset we used for our subsequent analyses.

### Simulations of the passaging experiment, DNA extraction, and sequencing

The simulation of our entire experiment, entails first a simulation of the viral populations and its diversification throughout 400 passages. This results in approximately 2×10^6^ virions with potentially different genotypes (although many virions carry identical genomes).

Subsequently, the simulated virions are sampled binomially according to the volume sampled from the wells in our experiments, which leads to a loss of rare genotypes and a distortion of their original frequencies. The sampled virions are then plated, their DNA is extracted, and, when simulating the ssDNA extraction protocol (see above), PCR amplified, or when simulating the dsDNA extraction protocol (see above), replicated in cells. The simulated plating, PCR amplification, and replication in cells can introduce errors into the genomes. Lastly, the simulated genomes are sequenced, which is implemented as binomial sampling and a worst case sequencing error rate.

For all the details on these simulations please consult the file “python-code/code/simulations.py” in the supplementary file “data+code-for-reviewers-Feb25-2026.zip”.

### Simulating sequences as expected under neutral evolution

To generate genotypes that would be expected under neutral evolution, we randomized the locations of mutations according to the following procedure.

First, we infer which of the mutations of a genotype have newly emerged during the passage experiment, rather than having been already present in the ancestral population. We do this by identifying the genotype in the ancestral population that has most mutations in common with the genotype in question. We assume this specific ancestral genotype to be the evolutionary origin, and consider all the mutations of the genotype not already present in the specific ancestor to have newly emerged. In case of multiple ancestral genotypes with the same number of shared mutations from the genotype in question, we assume the evolutionary origin to be the one that is most frequent in the ancestral population.

Second, we randomize the positions of these likely newly emerged mutations. Formally, if we have *n* newly emerged mutations, we sample without replacement n out of {1*, . . ., λ*}, where *λ* = 5386 is the length of the genome of φX174. (Thus, if, for example, a genotype has likely acquired a mutation at positions 1770 and 2966, this randomization could generate positions 3328 and 4599.) For the sake of simplicity, we randomize only the positions, rather than the positions and specific nucleotides substitutions. As a result, parallel evolution becomes more likely in these simulations, and the resulting randomization test is conservative, i.e. it underestimates the extent and significance of a deviation from neutrality.

Third, we adopted this procedure to generate viral populations with mutational patterns that would be expected under neutral evolution for each evolution line. Repeating this 10’000 times for all 31’229 distinct genotypes, we can estimate the probability of a mutated position being shared across multiple evolution lines, and compare it to the observed frequency (Figure 5A). In addition to considering single mutated positions, we also investigated combinations of mutated positions (Figure 5B–G). The clear conclusion from this analysis is that there is much more sharing of single mutated positions, and combinations of mutated positions than expected under neutral evolution (see main text). This could be due to a selective advantage that these mutations confer, or to a higher mutation probability at the positions that are shared across the evolution lines. Last, we investigated if combinations of mutations are shared more than expected given that already single mutations are shared more than expected under neutral evolution. To this end, we set the effective genome length to *λ* = 539 instead of *λ* = 5386. As a consequence, shared mutations become more likely. At this particular effective genome length, the observed number of shared mutated positions across four evolution lines is comparable to the neutral expectation (Figure 5A). The number of mutated positions shared across two or three evolution lines even exceeds this expectation, which makes the following analysis conservative.

In contrast to single mutated positions, the observed number of shared combinations of mutated positions is still much lower than the expectation at this low effective genome length (Figure 5B–G). This is strong evidence that these combinations have a selective advantage that exceeds the potential advantage that the individual mutations confer, i.e. interact under positive epistasis. Note that, while we cannot disentangle potential mutational bias from selective advantage for single mutations, once the effect of single mutations is subsumed by shortening the effective genome length, deviations from the neutral expectation must be due to positive epistatic interactions between the mutated sites.

### Linkage disequilibrium analysis to probe for genetic signatures of recombination

To investigate if recombination occurs to an extent that it leaves a detectable trace in the viral genomes, we performed a bioinformatic analysis of the sequencing data. Recent advanced methods to detect recombination are based on reconstructing phylogenies for multiple parts of the genome, for example GARD (Kosakovsky Pond et al., 2006). These, however, are challenging to apply because we have over 80’000 whole-genomes in our data set. For this reason, we specifically used older, more scalable methods that exploit patterns of genetic linkage that are indicative of recombination.

The well-known relationship between linkage disequilibrium and recombination rate by Hill and Robertson (1968) (Hartl and Clark, 2007, Hill and Robertson, 1968) cannot be used for our purpose because of some of the assumptions that go into the derivation of the relationship. The formula applies only if there are no new mutations (which occur during our experiment), if there is no selection (which we assume is acting), and, most importantly, when an evolutionary equilibrium has been reached (which is not the case in our experiments either).

We therefore used the even older formula for the decline of the linkage disequilibrium over time (Slatkin, 2008, Jennings, 1917), similar to recent methods developed by Romero and Feder (2024) for the estimation of effective recombination rates of HIV within the infected individual. From one generation to the next, the linkage disequilibrium between two mutations, *D* = *p*_12_ − *p*_1_ *p*_2_, declines by a factor (1 − *c*). Hereby *p*_1_, *p*_2_, and *p*_12_ denotes the frequencies of genotypes carrying mutation 1, 2, or both mutations, respectively, and *c* denotes the probability of recombination. The recombination rate is commonly assumed to scale with the distance between the mutations on the genome, and can hence be rewritten as *c* = *c*_1*bp*_ *d*, with *c*_1*bp*_ denoting the recombination rate at a genomic distance of a single base pair, and *d* denoting the genomic distance in base pairs.

Extrapolating this decline of *D* over time, we obtain the following relationship between the linkage disequilibrium *D*, the genomic distance of two mutations *d*, the recombination rate *c*_1*bp*_, and the generation *g* in our experiments:

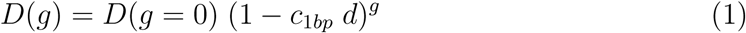

We fitted this relationship (Equation 1) to all the mutation pairs that have a non-zero *D* in the ancestor population. We normalized all measurements of *D* by dividing them by the non-zero *D* in the ancestor population. We obtain an estimate for *c*_1*bp*_ that is not significantly different from 0 (*c*_1*bp*_ = 0.0004 ± 0.0015). Thus, there is no signal for linkage disequilibrium decline over generations, and hence no evidence for recombination in our experiments. While this statistical test does not rule out any instance of recombination, it supports the assumption that recombination has not affected the evolutionary dynamics in a major way.

### Phylodynamics estimation of migration rates between the four evolution lines

To investigate to what extent the instances of parallel evolution that occur in our evolution experiment can bias the estimation of migration rates between the evolution lines, we conducted a phylodynamic analysis. As the evolution lines in our experiments evolve independently, i. e. there is no migration between these lines, the migration rates should be estimated to be zero.

In our experiments, we obtained a total of 80’886 whole-genome sequences that appeared more than 3 times in one of the samples. Of these sequences only 884 are distinct, i.e. many of the sequences are identical. This leads to the problem of polytomies in the reconstruction of the phylogenies, on which a phylodynamic analysis is built.

The high number of sequences and the polytomies make a one-step Bayesian estimation of the migration rates using the software BEAST (Drummond and Rambaut, 2007, Bouckaert et al., 2014, Vaughan et al., 2014, Bouckaert et al., 2019) prohibitive. We therefore resorted to a simpler, non-Bayesian two-step estimation procedure, ignoring the phylogenetic uncertainty inherent in the reconstructure of the tree. Technically, we conducted this analysis using *treetime* (Sagulenko et al., 2018), an efficient algorithm to generate timed phylogenetic trees from sequence data. Treetime allows a two-step *mugration* analysis (Lemey et al., 2009), which estimates migration rates from a phylogenetic tree analogously to mutation rates by adding a locus to the genomes that has its own mutation rate and designates the deme in which the sequences was sampled. The mutation rates between the, in our case, five demes then represent estimates of the migration rate between them.

To circumvent having to analyze all 80’886 sequences, we adopt the common strategy to subsample. We subsample 900 sequences, 100 from the ancestor population and another 100 from generation 196 and 412 of each of the four evolutiom lines, and repeat this analysis 100 times. This subsampling approach captures parts of the phylogenetic uncertainty that we did not incorporate directly.

Another complication that arises in the phylodynamic analysis of our experimental data is that the ancestor population, with which the four evolution lines were seeded, consists of multiple genotypes that arose through a common evolutionary history. This “panmictic past” will leave a signal for migration even in the independent lines. Our solution of this issue is to assign the ancestor its own deme, from which migration to each evolution line is possible. (Each evolution lines is also assigned a deme of its own.) This way, the common evolutionary history of the genotypes in the ancestor population can be separated from the potential migration of genotypes between evolution lines in the analysis (see Table S1).

### Code

The computer code developed and used for simulating phage evolution, analysing linkage disequilibrium, and conducting the migration rate estimation with treetime has been submitted as a zip-compressed supplementary file “data+code-for-reviewers-Feb25-2026.zip” along with this manuscript, and will be deposited at on zenodo upon acceptance.

## Supporting information

DNA extraction protocols developed and used in the study

File containing R-code and output of the analysis of the long-read sequencing data

## Acknowledgements

The authors thank James J. Bull for a decade of guidance and support of this study, Sylvain Gandon for his help designing the liquid titration of the bacteriophage in liquid, Marie-Agnes Petit, Claudia Igler, Silvia Kobbel, Léa Frachon for very fruitful discussions for the design of the DNA extraction protocols, Louis du Plessis for guidance on estimating migration rates with phylodynamic methods, and Claudia Igler and Carolin Wendling and Sira Ösze for detailed comments on the manuscript. RRR acknowledges the financial support of the Swiss National Science Foundation (grant number 31003A 179170).

## Supplementary Figures and Tables

**Figure S1:**
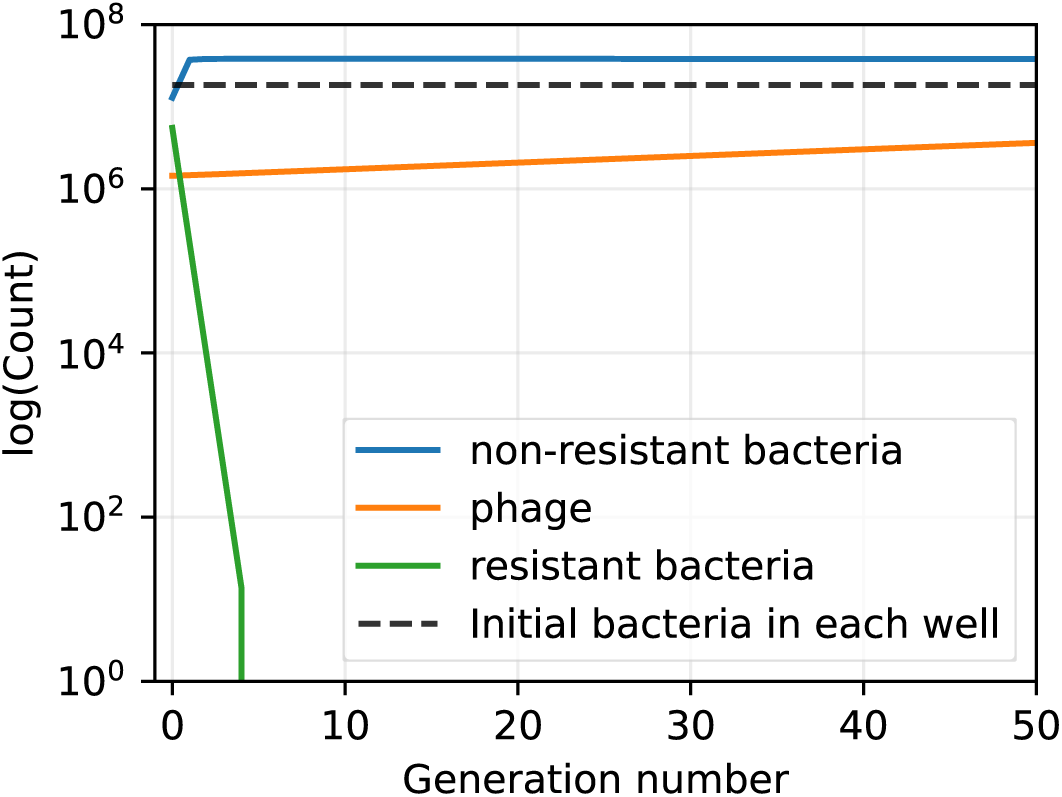
Population dynamic simulation showing that phage-resistant bacteria cannot evolve in our experimental setup. If bacteria evolve that are completely resistant to phage and carry no resistance cost, they are washed within a after a few transfers. This even applies to levels of resistant bacteria that are unrealistically high, such as the hypothetical level of 30% used in the simulation shown: they are washed out within 4 transfers. This is because in our experiment we transfer 2% of the phages (and some bacteria) every 26 minutes, (which corresponds to their lysis time) and transfer them into a well that has been seeded with a billion bacteria from a stock. The python code of this simulation can be found in the file “python-code/code/simulation-showing-the-impossibility-of-bacterial-phage-resistance.ipynb” in the supplementary zip-archive “data+code-for-reviewers-Feb25-2026.zip”.

**Figure S2:**
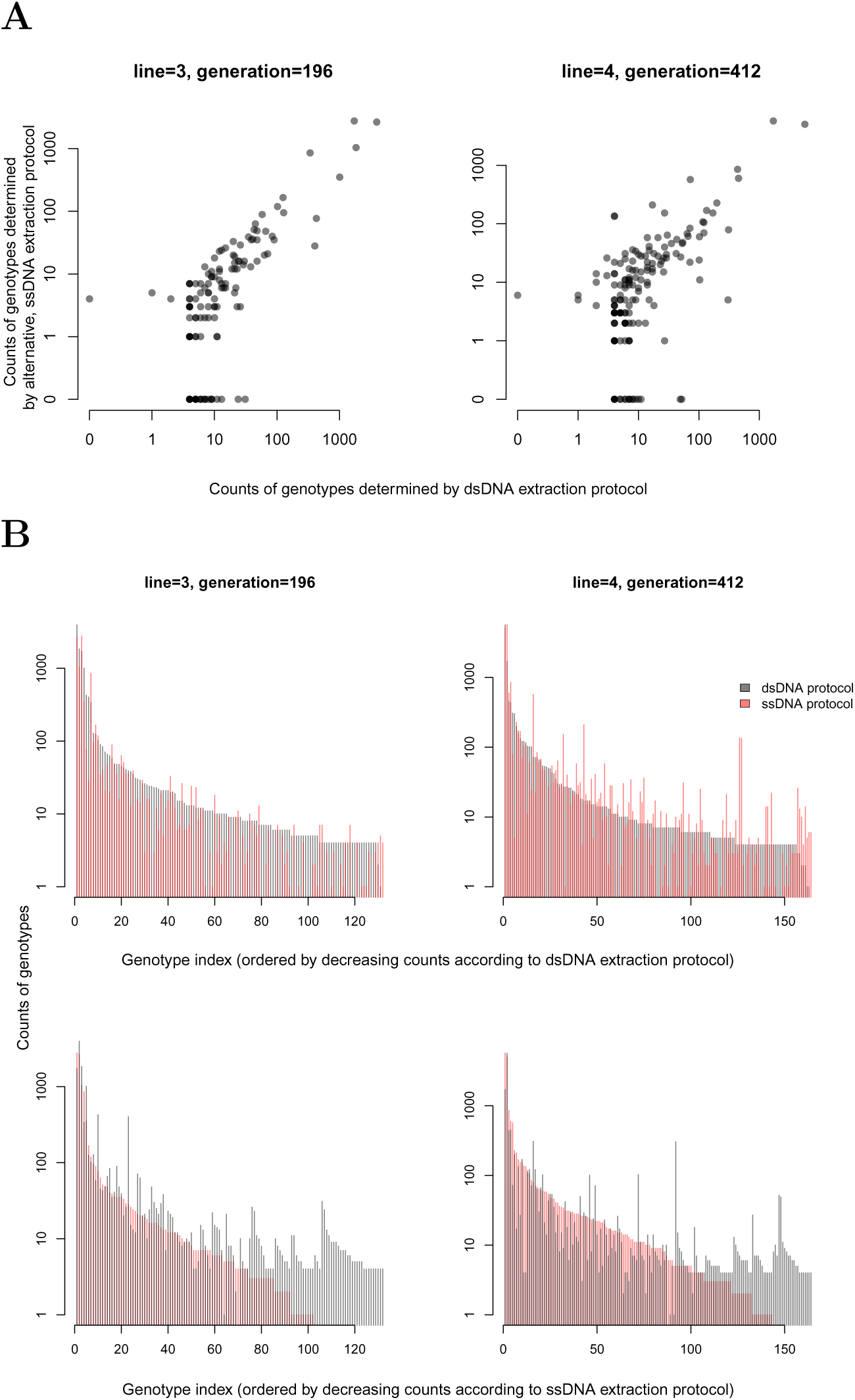
Comparison of the dsDNA extraction protocol and the alternative ssDNA extraction protocol in 2 of our 9 samples. **(A)** Comparison of the frequencies of the genotypes. **(B)** Comparison of the genotype counts distributions. The top and bottom rows display the genotype counts order by decreasing counts according to the dsDNA and ssDNA extraction protocol, respectively. The dsDNA extraction protocol was more efficient as it generates more distinct genotypes (see the file “R-code/code/stats-for-paper-rrr.html” in the supplementary zip-archive “data+code-for-reviewers-Feb25-2026.zip”).

**Figure S3:**
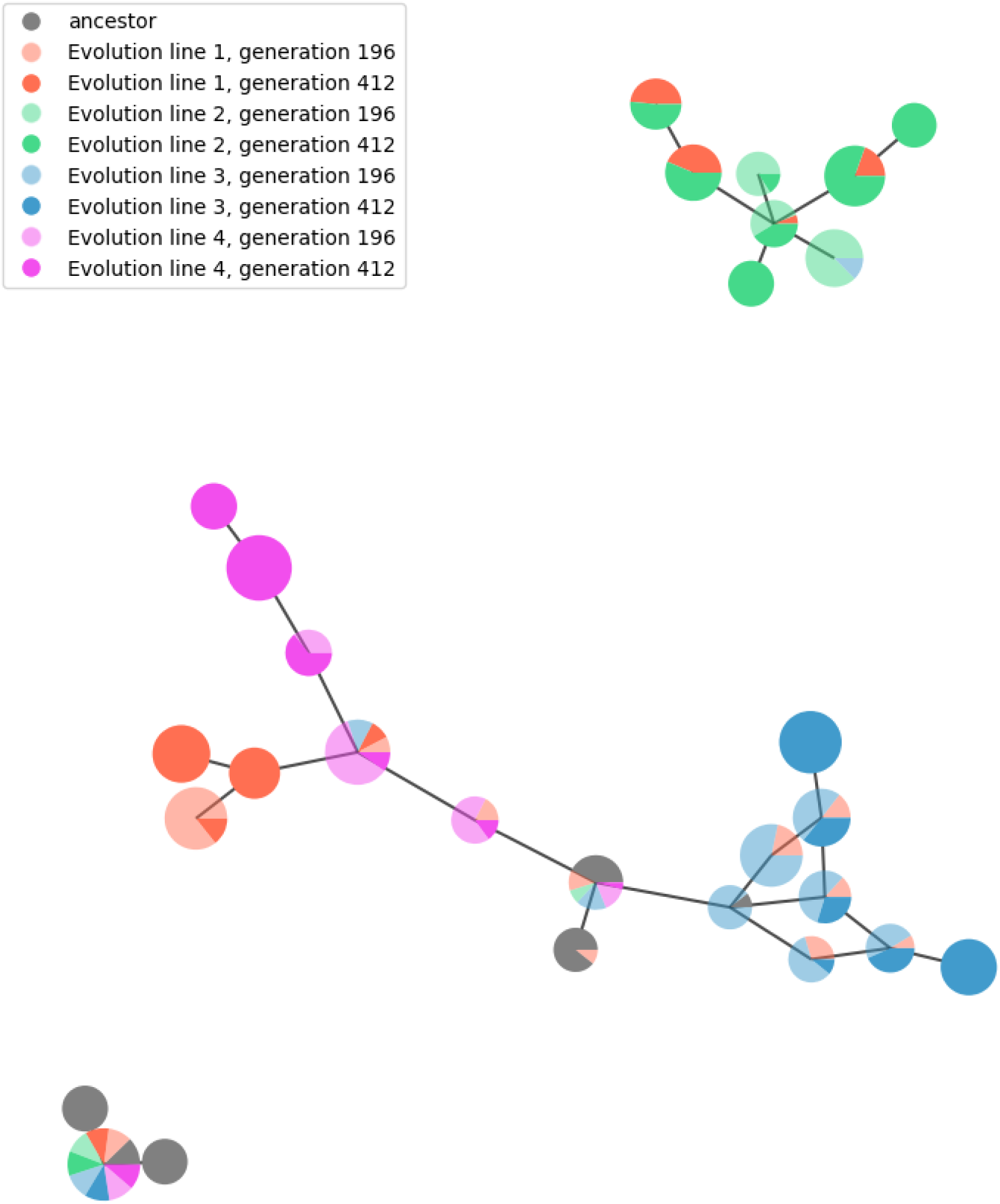
Genotype network graph of the 50 most frequent genotypes out of the 884 that we observed in our experiment.

**Table S1:** Migration rate estimate from each deme to each other deme (average over estimates for the 100 subsamples). Deme A is the ancestor population, demes EL1–4 are evolution lines 1–4. The direction of migration goes from column to row. The estimated rates of migration from the ancestor population to the evolution lines can be seen in the first column of the matrix, and has consistently high values. This is consistent with the fact that each evolution line has been seeded with the ancestor population, i. e. migration actually happened in the experiment. The other four columns show estimates of migration between the evolution lines, and from the evolution lines back to the ancestor. These migration rates are zero in the experiment, but are wrongly estimated to be non-zero. This bias arises because of parallel evolution between the evolution lines. Between evolution lines 1 and 2, for example, there are particularly many genotypes that evolved in parallel by generation 412 (see Figure 4), leading to the largest migration rate estimates between these lines (0.00168 and 0.00182). The basis to calculate the overall migration rate estimate between the lines were columns 2–5 in the migration rate estimate matrices. Leaving out the first column in calculating the overall migration rate ensures that the migration rate is not confounded by the “panmictic past” of the ancestor population.

|  | A | EL1 | EL2 | EL3 | EL4 |
| --- | --- | --- | --- | --- | --- |
| A | 0.00000 | 0.00021 | 0.00020 | 0.00022 | 0.00019 |
| EL1 | 0.00153 | 0.00000 | 0.00182 | 0.00095 | 0.00098 |
| EL2 | 0.00138 | 0.00168 | 0.00000 | 0.00065 | 0.00059 |
| EL3 | 0.00132 | 0.00077 | 0.00056 | 0.00000 | 0.00062 |
| EL4 | 0.00145 | 0.00104 | 0.00069 | 0.00084 | 0.00000 |

## References

Acevedo A, Brodsky L & Andino R (2014). Mutational and fitness landscapes of an RNA virus revealed through population sequencing. Nature 505(7485): 686–690.

Baker CW, Miller CR, Thaweethai T, Yuan J, Baker MH, Joyce P & Weinreich DM (2016). Genetically Determined Variation in Lysis Time Variance in the Bacteriophage *ϕX*174. G3 (Bethesda) 6(4):939–55.

Bedford T, Cobey S & Pascual M (2011). Strength and tempo of selection revealed in viral gene genealogies. BMC evolutionary biology 11:220.

Bertels F, Leemann C, Metzner KJ & Regoes R (2019). Parallel evolution of HIV-1 in a long-term experiment. Mol. Biol. Evol. 36(11):2400–2414.

Bons E, Leemann C, Metzner KJ & Regoes RR (2020). Long-term experimental evolution of HIV-1 reveals effects of environment and mutational history. PLoS Biol 18(12):e3001010.

Bons E & Regoes RR (2018). Virus dynamics and phyloanatomy: Merging population dynamic and phylogenetic approaches. Immunol. Rev. 285(1):134–146.

Bouckaert R, Heled J, Kühnert D, Vaughan T, Wu CH, Xie D, Suchard MA, Rambaut A & Drummond AJ (2014). BEAST 2: a software platform for Bayesian evolutionary analysis. PLoS Comput. Biol. 10(4):e1003537.

Bouckaert R, Vaughan TG, Barido-Sottani J, Duchêne S, Fourment M, Gavryushkina A, Heled J, Jones G, Kühnert D, De Maio N & others (2019). Beast 2.5: An advanced software platform for bayesian evolutionary analysis. PLoS computational biology 15(4):e1006650.

Bull JJ, Badgett MR, Wichman HA, Huelsenbeck JP, Hillis DM, Gulati A, Ho C & Molineux IJ (1997). Exceptional convergent evolution in a virus. Genetics 147(4):1497–507.

Cuevas JM, Duffy S & Sanjuán R (2009). Point mutation rate of bacteriophage PhiX174. Genetics 183(2):747–9.

Denhardt DT & Silver RB (1966). An analysis of the clone size distribution of *φ*x174 mutants and recombinants. Virology 30(1):10–19.

Desai MM & Fisher DS (2007). Beneficial mutation–selection balance and the effect of linkage on positive selection. Genetics 176(3):1759–1798.

Domingo-Calap P, Cuevas JM & Sanjuán R (2009). The fitness effects of random mutations in single-stranded DNA and RNA bacteriophages. PLoS Genet. 5(11):e1000742.

Drummond AJ & Rambaut A (2007). BEAST: Bayesian evolutionary analysis by sampling trees. BMC Evol. Biol. 7:214.

Düx A, Lequime S, Patrono LV, Vrancken B, Boral S, Gogarten JF, Hilbig A, Horst D, Merkel K, Prepoint B & others (2020). Measles virus and rinderpest virus divergence dated to the sixth century bce. Science 368 (6497):1367–1370.

Eid J, Fehr A, Gray J, Luong K, Lyle J, Otto G, Peluso P, Rank D, Baybayan P, Bettman B & others (2009). Real-time dna sequencing from single polymerase molecules. Science 323(5910):133–138.

Faria NR, Rambaut A, Suchard MA, Baele G, Bedford T, Ward MJ, Tatem AJ, Sousa JaD, Arinaminpathy N, Pépin J, Posada D, Peeters M, Pybus OG & Lemey P (2014). HIV epidemiology. The early spread and epidemic ignition of HIV-1 in human populations. Science 346(6205):56–61.

Fraser C, Donnelly CA, Cauchemez S, Hanage WP, Van Kerkhove MD, Hollingsworth TD, Griffin J, Baggaley RF, Jenkins HE, Lyons EJ, Jombart T, Hinsley WR, Grassly NC, Balloux F, Ghani AC, Ferguson NM, Rambaut A, Pybus OG, Lopez-Gatell H, Alpuche-Aranda CM, Chapela IB, Zavala EP, Guevara DME, Checchi F, Garcia E, Hugonnet S & Roth C (2009). Pandemic potential of a strain of influenza A (H1N1): early findings. Science 324(5934):1557–61.

Frost SD, Pybus OG, Gog JR, Viboud C, Bonhoeffer S & Bedford T (2015). Eight challenges in phylodynamic inference. Epidemics 10:88–92.

Gerrish PJ & Lenski RE (1998). The fate of competing beneficial mutations in an asexual population. Genetica 102:127–144.

Gire SK, Goba A, Andersen KG, Sealfon RSG, Park DJ, Kanneh L, Jalloh S, Momoh M, Fullah M, Dudas G, Wohl S, Moses LM, Yozwiak NL, Winnicki S, Matranga CB, Malboeuf CM, Qu J, Gladden AD, Schaffner SF, Yang X, Jiang PP, Nekoui M, Colubri A, Coomber MR, Fonnie M, Moigboi A, Gbakie M, Kamara FK, Tucker V, Konuwa E, Saffa S, Sellu J, Jalloh AA, Kovoma A, Koninga J, Mustapha I, Kargbo K, Foday M, Yillah M, Kanneh F, Robert W, Massally JLB, Chapman SB, Bochicchio J, Murphy C, Nusbaum C, Young S, Birren BW, Grant DS, Scheiffelin JS, Lander ES, Happi C, Gevao SM, Gnirke A, Rambaut A, Garry RF, Khan SH & Sabeti PC (2014). Genomic surveillance elucidates Ebola virus origin and transmission during the 2014 outbreak. Science 345(6202):1369–72.

Hartl D & Clark A. Principles of Population Genetics. Sinauer, 2007. ISBN 9780878933082.

Hill W & Robertson A (1968). Linkage disequilibrium in finite populations. Theoretical and applied genetics 38:226–231.

Hillis DM, Bull JJ, White ME, Badgett MR & Molineux IJ (1992). Experimental phylogenetics: generation of a known phylogeny. Science 255(5044): 589–92.

Holmes EC. The evolution and emergence of RNA viruses. Oxford University Press, 2009.

Jennings HS (1917). The numerical results of diverse systems of breeding, with respect to two pairs of characters, linked or independent, with special relation to the effects of linkage. Genetics 2(2):97.

Keele BF, Giorgi EE, Salazar-Gonzalez JF, Decker JM, Pham KT, Salazar MG, Sun C, Grayson T, Wang S, Li H, Wei X, Jiang C, Kirchherr JL, Gao F, Anderson JA, Ping LH, Swanstrom R, Tomaras GD, Blattner WA, Goepfert PA, Kilby JM, Saag MS, Delwart EL, Busch MP, Cohen MS, Montefiori DC, Haynes BF, Gaschen B, Athreya GS, Lee HY, Wood N, Seoighe C, Perelson AS, Bhattacharya T, Korber BT, Hahn BH & Shaw GM (2008). Identification and characterization of transmitted and early founder virus envelopes in primary HIV-1 infection. Proc Natl Acad Sci U S A 105(21):7552–7.

Koelle K & Rasmussen DA (2025). Phylodynamics beyond neutrality: the impact of incomplete purifying selection on viral phylogenies and inference. Philosophical Transactions B 380(1919):20230314.

Kosakovsky Pond SL, Posada D, Gravenor MB, Woelk CH & Frost SD (2006). Automated phylogenetic detection of recombination using a genetic algorithm. Molecular biology and evolution 23(10):1891–1901.

Lemey P, Rambaut A, Drummond AJ & Suchard MA (2009). Bayesian phylogeography finds its roots. PLoS Comput. Biol. 5(9):e1000520.

Li H, Bar KJ, Wang S, Decker JM, Chen Y, Sun C, Salazar-Gonzalez JF, Salazar MG, Learn GH, Morgan CJ, Schumacher JE, Hraber P, Giorgi EE, Bhattacharya T, Korber BT, Perelson AS, Eron JJ, Cohen MS, Hicks CB, Haynes BF, Markowitz M, Keele BF, Hahn BH & Shaw GM (2010). High Multiplicity Infection by HIV-1 in Men Who Have Sex with Men. PLoS Pathog. 6(5):e1000890.

Lorenzo-Redondo R, Fryer HR, Bedford T, Kim EY, Archer J, Kosakovsky Pond SL, Chung YS, Penugonda S, Chipman JG, Fletcher CV, Schacker TW, Malim MH, Rambaut A, Haase AT, McLean AR & Wolinsky SM (2016). Persistent HIV-1 replication maintains the tissue reservoir during therapy. Nature 530(7588):51–6.

Miller CR, Nagel AC, Scott L, Settles M, Joyce P & Wichman HA (2016). Love the one you’re with: replicate viral adaptations converge on the same phenotypic change. PeerJ 4:e2227.

Müller NF, Kistler KE & Bedford T (2022). A bayesian approach to infer recombination patterns in coronaviruses. Nature communications 13(1):4186.

Nadeau SA, Vaughan TG, Scire J, Huisman JS & Stadler T (2021). The origin and early spread of SARS-CoV-2 in europe. Proc. Natl. Acad. Sci. U. S. A. 118(9):e2012008118.

Neher RA & Leitner T (2010). Recombination rate and selection strength in hiv intra-patient evolution. PLoS computational biology 6(1):e1000660.

Ochsner N, Bouman J, Vaughan T, Stadler T, Bonhoeffer S & Regoes R (2026). Viral simulation reveals overestimation bias in within-host phylodynamic migration rate estimates under selection. Molecular biology and evolution 43(2).

Posada D & Crandall KA (2002). The effect of recombination on the accuracy of phylogeny estimation. Journal of molecular evolution 54(3):396–402.

Prosperi MCF & Salemi M (2012). QuRe: software for viral quasispecies reconstruction from next-generation sequencing data. Bioinformatics 28(1): 132–3.

Rasmussen DA & Stadler T (2019). Coupling adaptive molecular evolution to phylodynamics using fitness-dependent birth-death models. eLife 8.

Ratmann O, Hodcroft EB, Pickles M, Cori A, Hall M, Lycett S, Colijn C, Dearlove B, Didelot X, Frost S, Hossain ASMM, Joy JB, Kendall M, Kühnert D, Leventhal GE, Liang R, Plazzotta G, Poon AFY, Rasmussen DA, Stadler T, Volz E, Weis C, Leigh Brown AJ & Fraser C (2017). Phylogenetic Tools for Generalized HIV-1 Epidemics: Findings from the PANGEA-HIV Methods Comparison. Mol. Biol. Evol. 34(1):185–203.

Romero EV & Feder AF (2024). Elevated hiv viral load is associated with higher recombination rate in vivo. Molecular Biology and Evolution 41(1): msad260.

Sagulenko P, Puller V & Neher RA (2018). Treetime: Maximum-likelihood phylodynamic analysis. Virus evolution 4(1):vex042.

Salemi M & Rife B (2016). Phylogenetics and Phyloanatomy of HIV/SIV Intra-Host Compartments and Reservoirs: The Key Role of the Central Nervous System. Curr. HIV Res. 14(2):110–20.

Sanger F, Air GM, Barrell BG, Brown NL, Coulson AR, Fiddes CA, Hutchison CA, Slocombe PM & Smith M (1977). Nucleotide sequence of bacteriophage phi X174 DNA. Nature 265(5596):687–95.

Schlub TE, Grimm AJ, Smyth RP, Cromer D, Chopra A, Mallal S, Venturi V, Waugh C, Mak J & Davenport MP (2014). Fifteen to twenty percent of hiv substitution mutations are associated with recombination. Journal of virology 88(7):3837–3849.

Sinsheimer RL, Starman B, Nagler C & Guthrie S (1962). The process of infection with bacteriophage *φ*x174: I. evidence for a “replicative form”. Journal of molecular biology 4(3):142–160.

Slatkin M (2008). Linkage disequilibrium—understanding the evolutionary past and mapping the medical future. Nature Reviews Genetics 9(6):477–485.

Turner PE, McBride RC, Duffy S, Montville R, Wang LS, Yang YW, Lee SJ & Kim J (2012). Evolutionary genomics of host-use in bifurcating demes of rna virus phi-6. BMC evolutionary biology 12:153.

Vaughan TG, Kühnert D, Popinga A, Welch D & Drummond AJ (2014). Efficient Bayesian inference under the structured coalescent. Bioinformatics 30(16):2272–9.

Wenger AM, Peluso P, Rowell WJ, Chang PC, Hall RJ, Concepcion GT, Ebler J, Fungtammasan A, Kolesnikov A, Olson ND & others (2019). Accurate circular consensus long-read sequencing improves variant detection and assembly of a human genome. Nature Biotechnology 37(10):1155–1162.

Wichman HA, Badgett MR, Scott LA, Boulianne CM & Bull JJ (1999). Different trajectories of parallel evolution during viral adaptation. Science 285(5426):422–4.

Wichman HA, Millstein J & Bull JJ (2005). Adaptive molecular evolution for 13,000 phage generations: a possible arms race. Genetics 170(1):19–31.

Zagordi O, Bhattacharya A, Eriksson N & Beerenwinkel N (2011). ShoRAH: estimating the genetic diversity of a mixed sample from next-generation sequencing data. BMC Bioinformatics 12:119.

