## Supplementary material for "Parallel evolution of full-length genomes in a long-term evolution experiment with phage φX174": DNA extraction protocols developed and used in the study

#### 1 phiX174 dsDNA extraction protocol

This protocol has been designed to extract phiX174 dsDNA for PacBio sequencing, without having a lot of phage recombination. It results in around 1 microgramm of dsDNA with high purity (whole genome linear dsDNA, A260/A280 = 1.8, A260/A230 >2, no DNA and no RNA from the bacteria). Inspired from `http://wiki.bx.psu.edu/PhiX:DsDNA_isolation`.

##### 1.1 Reagents

- LB Miller
- $MgCl_2$ 1M
- $CaCl_2$ 1M
- Petri dishes poured with Hard Agar (LB + 10 mM $MgCl_2$ + 2 mM $CaCl_2$ + 1.5% Agar)
- Soft Agar (LB + 10 mM $MgCl_2$ + 2 mM $CaCl_2$ + 0.5% Agar)
- 0.22 $\mu$m filters
- Chloramphenicol 100 mg/mL
- New Englands Biolabs Monarch Plasmid Miniprep kit
- Nuclease free water
- New Englands Biolabs Cutsmart buffer
- New Englands Biolabs Pst1-HF
- Ampure beads
- Qubit BR dsDNA kit
- Agilent Tape Station genomic kit

##### 1.2 Day 1: Phage amplification

1. Grow E. coli C in LB until stationary phase at 37°C.
2. Take 400 $\mu$L of fresh stationary phase E. coli C, add 20 µL of phage lysate, 6 mL of soft agar, $CaCl_2$ at a final concentration of 2 mM and $MgCl_2$ at a final concentration of 10 mM. Mix and pour on a Hard Agar plate.
3. Incubate overnight at 37°C.
4. Start an E. coli C culture in LB that you grow overnight at 37°C with aeration.

##### 1.3 Day 2 and 3: dsDNA amplification, plasmid extraction, DNA cleaning and DNA quality assessment

1. Inoculate 200 mL of LB with 2 mL of E. coli C in stationary phase. Incubate at 37°C for exactly 1h30 with aeration.
2. During incubation, extract phages by ading 3 mL of LB, mixing and then pipetting the LB into eppendorf tubes. You should get ≈ 2 mL of phage lysate. Then, centrifuge for 5 minutes at maximum speed and filter these solutions using 0.22 $\mu$m filters. Store in the fridge.
3. After 1h30 of bacterial growth, add $MgCl_2$ at 10 mM (2 mL) and $CaCl_2$ at 2 mM (400 $\mu$L). Add 2 mL of the phage lysate and incubate at 37°C for exactly 5 minutes.
4. Add 600 $\mu$L of chloramphenicol at 100 mg/mL and incubate for 3 hours at 37°C with aeration.
5. Split the 200 mL cultures into 4 50-mL tubes. Centrifuge at maximal speed for 5 minutes.
6. Discard supernatant. Resuspend each pellet in NEB resuspension buffer (from NEB Monarch Plasmid miniprep kit).
7. Do one tube at a time. For each tube, add 200 microlitres of NEB lysis buffer. Carefully invert the tube once or twice and directly add 400 $\mu$L of NEB neutralisation buffer. Be careful not to delay the neutralisation. Mix very carefully by inverting the tubes until the solution becomes yellow. You can keep it on the bench while you are preparing the remaining tubes.
8. Centrifuge for 5 minutes at 16 000 g.
9. Add one supernatant to one NEB column. Centrifuge for 1 minute at 16 000 g. Discard flowthrough.
10. Add the second supernatant to the same column. Centrifuge for 1 minute at 16 000 g. Discard flowthrough.
11. Add the third supernatant to the same column. Centrifuge for 1 minute at 16 000 g. Discard flowthrough.
12. Add the fourth supernatant to the same column. Centrifuge for 1 minute at 16 000 g. Discard flowthrough.
13. Add 200 $\mu$L of NEB Wash solution 1. Centrifuge for 1 minute at 16 000 g. Discard flowthrough.
14. Add 400 $\mu$L of NEB Wash solution 2. Centrifuge for 1 minute at 16 000 g.
15. Discard flowthrough without touching the column. Add 30 $\mu$L of nuclease free water at the center of the column. Let rest for 1 minute. Centrifuge at 16 000 g for 1 minute.
16. If needed, samples can be kept in the fridge.
17. Open the phage DNA by adding 10% of NEB Cutsmart buffer and 1 $\mu$L of Pst1-HF. Incubate 10 minutes at 37°C.
18. Clean the phage DNA by using Ampure beads as in the ssDNA extraction protocol.
19. Quantify DNA using Qbit (dsDNA BR Kit) and purity using Nanodrop according to manufacturer instructions. Nanodrop and Qubit concentration will be very different (usually by a factor ≈ 10). We were not able to find the source of this discrepancy, nor to remove it. However, in our case, this did not pose any problem for PacBio sequencing.
20. Run an electrophoresis using the Agilent tape station according to manufacturer instructions.
21. Samples can be kept in the fridge.

### 2 phiX174 ssDNA extraction protocol

Objective: setting up a DNA extraction of E. coli C phage phiX174 DNA (5.3 kb circular ssDNA genome), that is of good quality for PacBio sequencing (whole genome linear dsDNA, A260/A280 = 1.8, A260/A230 >2, no DNA and no RNA from the bacteria, in high amount (at the very minimum 500 ng per sample). Inspired from:

- http://www.protocol-online.org/biology-forums/posts/40049.html
- http://wiki.bx.psu.edu/PhiX:SsDNA_isolation
- Mayjonade et al. Mayjonade et al. (2016)
- Schalamun et al. Schalamun and Schwessinger (2017)

#### 2.1 Reagents

- LB Miller
- Hard Agar (LB Miller + 15 g/L Agar + 2 mM $CaCl_2$ + 10 mM $MgCl_2$) and Soft Agar (LB Miller + 5 g/L Agar + 2 mM $CaCl_2$ + 10 mM $MgCl_2$)
- filter 0.22 $\mu$m
- Nuclease free water
- $MgCl_2$ 1M
- $CaCl_2$ 1M
- PEG 8000 30% 1.5M NaCl
- DNase buffer from NEB
- DNase 1
- RNase A from ThermoFischer
- EDTA 50mM,pH8
- SDS 20%
- Pronase
- NaCl 5M
- 100% Ethanol at -20°C.
- 70% Ethanol at -20°C.
- Q5 polymerase and buffer
- dNTP 10mM
- Primer PhiX1F 10 mM (ATCGCTTCCATGACGCAGAA)
- Primer PhiX1R 10 mM (TGCAGGTTGGATACGCCAAT)
- 100% DMSO
- Agilent Tape Station genomic kit
- Ampure Beads
- Magnetic rack
- Absolute ethanol (for DNA extraction) at room temperature
- EB Buffer (Quiagen).
- dsDNA BR Qubit Kit

#### 2.2 Day 1 and 2: Phage amplification and PEG precipitation

1. Take 400 $\mu$L of fresh stationary phase E. coli C, add 20 µL of phage lysate, 6 mL of soft agar, $CaCl_2$ at a final concentration of 2 mM and $MgCl_2$ at a final concentration of 10 mM. Mix and pour on a Hard Agar plate. Incubate overnight at 37°C.
2. On the soft agar, add 3 mL of LB broth, shake, mix well. Then pipette 2 mL again and filter them with a 0.22 $\mu$m filter. Keep the flowthrough. You will need 1 mL for the following steps. This can be kept in the fridge for some time.
3. For 1 mL of phages, add 500 $\mu$L of PEG 30%, NaCl 1.5M. Mix well.
4. Leave in the fridge for 24 hours.

#### 2.3 Day 3: Extraction and precipitation of phiX174 ssDNA

1. Unfreeze the DNase buffer. Take the pronase out of the freezer and let it gently warm. Preheat three heaters, one at 37°C, a second one at 75°C and a third one at 40°C.
2. Take out the PEG precipitation from the fridge and centrifuge it 10 minutes at 13 000 g at 4°C.
3. Discard 1 mL of the supernatant with a blue tip and centrifuge the tube again for 2 minutes at 13 000 g at 4°C.
4. Discard the supernatant with a yellow tip and resuspend in 200 $\mu$L of DNase A buffer.
5. Leave on ice for 10 minutes.
6. Add 2 $\mu$L of DNase 1 and RNase A and incubate at 37°C for 10 minutes.
7. Add 20 $\mu$L of EDTA, mix and incubate for 10 minutes at 75°C.
8. Add 5 $\mu$L of SDS 20% and 5 $\mu$L of pronase. Incubate for 2.5 hours at 40°C.
9. Estimate precisely the total volume of your sample. Add 10% of its volume of NaCl 5M and 2 volumes of icecold 100% pure ethanol. Put in the freezer overnight.

#### 2.4 Day 4: DNA amplification

1. Cool down the centrifuge at 0°C. Unfreeze DMSO gently.
2. Centrifuge your sample at max speed for 10 minutes at 0°C.
3. Carefully removed supernatant, add 200 $mu$L of 70% ice cold ethanol and centrifuge at max speed at 0°C for 2 minutes.
4. Remove supernatant carefully, leave on the bench for 15 minutes (to evaporate the remaining ethanol) and resuspend in 100 $\mu$L of nuclease free water. Assess the quantity and purity of the DNA using a spectrophotometer. I usually get very good A260/A280 ratios, variables A260/A230 ratios (it does not seem to be a problem) and DNA concentrations between 5 and 20 $\mu$g/mL. You can't keep this DNA for long, do not stock before doing the PCR.
5. The PCR is tricky and tend to randomly fail. I have not found any way to predict whether it will work or not. Moreover, to decrease PCR bias, multiple PCR will be run for the same samples. Make a PCR master mix for 10*n samples (ideally, add a positive control), following the composition per tube in the next table. Add polymerase at the last moment. Keep polymerase at -20°C. Dispatch in tubes.

| Component | Volume | Final Concentration |
|---|---|---|
| Nuclease Free Water | 21.5 $\mu$L | |
| 5X Q5 Reaction Buffer | 10 $\mu$L | 1X |
| DMSO 100% | 2 $\mu$L | 4% |
| 10 $\mu$M Forward Primer | 2.5 $\mu$L | 0.5 $\mu$M |
| 10 $\mu$M Reverse Primer | 2.5 $\mu$L | 0.5 $\mu$M |
| 10mM dNTP | 1 $\mu$L | 200 $\mu$M |
| Q5 Polymerase | 0.5 $\mu$L | 0.02 U/$\mu$L |
| Phage DNA | 10 $\mu$L | $<$ 1000 ng |

6. Create the following program on the thermocycler. Preheat the thermocycler.

| Step | Temperature | Time |
|---|---|---|
| Initial Denaturation | 98°C | 30 seconds |
| Denaturation x32 | 98°C | 10 seconds |
| Anhealing x32 | 68°C | 30 seconds |
| Elongation x32 | 72°C | 108 seconds |
| Final extension | 72°C | 2 minutes |
| Storage | 4°C | indefinite |

#### 2.5 Day 5: DNA cleaning and quality assessment

1. Check the quality of the PCR amplification by running an electrophoresis with Agilent Tape Station (genomic kit). Follow the manufacturer instructions. You want to make sure that you just have a band at 5.3 kb and especially no amplification at 2.5 kb. If this is the case, continue the protocol. If not, redo the PCR.
2. Before starting, preheat the EB Buffer at 50°C, let the beads reach room temperature, make fresh wash solution (70% Ethanol). To avoid breaking the DNA, be gentle when pipetting.

3. Mix all the PCR products from the same sample in a single 2 mL tube and add 0.8 volume of Ampure beads (make sure the beads are completely resuspended before use). Mix well.
4. Incubate with a gentle agitation for 10 minutes at room temperature.
5. Spin down the tube. Place the tube in a magnetic rack for until the solution becomes clear (usually a couple of minutes is enough).
6. Remove the supernatant without disturbing the beads pellet.
7. Add 1 mL of wash solution.
8. Remove the supernatant and add 1 mL of wash solution.
9. Remove the supernatant and add 1 mL of wash solution.
10. Remove the supernatant. Spin down and remove the remaining wash solution. Do not let any trace of ethanol and do not let the beads dry. Add 80 $\mu$L of EB buffer preheated at 50°C.
11. Resuspend the beads and incubate at least 15 minutes at 37°C.
12. Place the tube in the magnetic rack. Let the solution become clear and pipette 75$\mu$ L of it in a new tube.
13. Assess the purity of the DNA using a nanodrop according to manufacturer instructions.
14. Assess DNA concentration using Qubit measurements according to manufacturer instructions (BR dsDNA kit).
15. If needed, keep DNA in the fridge.

#### References

Mayjonade B, Gouzy J, Donnadieu C, Pouilly N, Marande W, Callot C, Langlade N & Muños S (2016). **Extraction of high-molecular-weight genomic DNA for long-read sequencing of single molecules**. Biotechniques **61**(4):203–205.

Schalamun M & Schwessinger B (2017). **DNA size selection (>1kb) and clean up using an optimized SPRI beads mixture**. protocols.io.
