## Supplementary material for "Parallel evolution of full-length genomes in a long-term evolution experiment with phage φX174": File containing R-code and output of the analysis of the long-read sequencing data: stats-for-paper-rrr.html

Extracting statistics from the whole-population whole-genome phiX174 sequencing data


Code 

- Show All Code
- Hide All Code

### Extracting statistics from the whole-population whole-genome phiX174 sequencing data

###### Roland R. Regoes

###### February 19, 2026

### Read data

```
hpls <- read.table("../../python-code/data/haplotypes_run2.csv", sep=",", header=T)
barcds <- read.table("../../python-code/data/run2_barcodes.csv",
                     colClasses=c("character","numeric","numeric","numeric","character"),
                     sep=",", header=T)
n.reads <- sum(hpls[,-1])
attr(n.reads, "definition") <- "The total number of reads extracted from data in haplotypes_run2.csv"
n.reads.new <- sum(hpls[,2:10])
attr(n.reads.new, "definition") <- "The total number of reads extracted from data in haplotypes_run2.csv DISREGARDING THE OLD PROTOCOLL IN COLUMNS X16 AND X17"
```

Please note that we use the term “haplotype” in many file, variable
and function names in the code synonymously to the term “genotype” that
we use in the paper.

### Statistics on number of genotypes

**How many genotypes do we have in the raw data?**

```
sum(hpls[,-1])  # 150414
```

```
## [1] 150414
```

**How many, if we exclude reads generated by DNA extraction
protocol 2?**

```
sum(hpls[,2:10])  # 112734
```

```
## [1] 112734
```

**How many reads do we have per sample, if we exclude reads
generated by DNA extraction protocol 2?**

```
colSums(hpls[,2:10])
```

```
##   X01   X02   X03   X08   X09   X10   X11   X12   X15 
##  9169  9888 11125 16522 14001  8866 13088 13429 16646
```

```
#  X01   X02   X03   X08   X09   X10   X11   X12   X15 
# 9169  9888 11125 16522 14001  8866 13088 13429 16646 
colSums(hpls[,11:12])  # protocol 2
```

```
##   X16   X17 
## 13286 24394
```

```
#  X16   X17 
#13286 24394
```

**How many unique genotypes do we have in the raw
data?**

```
str(hpls)
```

```
## 'data.frame':    41161 obs. of  12 variables:
##  $ haplotype: chr  "" "1300:A->G;1303:C->G;1426:G->A;1638:T->G" "4630:C->T" "3339:G->A" ...
##  $ X01      : num  3138 1294 427 411 343 ...
##  $ X02      : num  1083 8 0 0 2 ...
##  $ X03      : num  1222 4 0 0 0 ...
##  $ X08      : num  1723 21 0 0 1 ...
##  $ X09      : num  1639 12 0 0 0 ...
##  $ X10      : num  974 0 0 0 0 0 4 0 0 1 ...
##  $ X11      : num  1340 0 0 0 0 0 1 0 0 0 ...
##  $ X12      : num  1460 0 0 0 0 2 2 0 0 6 ...
##  $ X15      : num  1704 2 0 0 0 ...
##  $ X16      : num  2769 19 0 0 5 ...
##  $ X17      : num  5728 0 0 0 0 ...
```

```
sum(rowSums(hpls[,-1])>0)    # including samples generated with protocol 2: 41161
```

```
## [1] 41161
```

```
sum(rowSums(hpls[,2:10])>0)  # without protocol 2: 31229
```

```
## [1] 31229
```

**How many unique genotypes do we have when restricting data to
>3 reads per sample?**

```
hpls.gt3.too.lenient <- hpls[rowSums(hpls[,-1])>3,]
ind <- sapply(hpls.gt3.too.lenient$haplotype,
              function(h)
                  any(hpls.gt3.too.lenient[hpls.gt3.too.lenient$haplotype==h,2:10]>3))
hpls.gt3 <- hpls.gt3.too.lenient[ind,]
rm(ind)
dim(hpls.gt3)[[1]]  # 884
```

```
## [1] 884
```

**What total (i.e. not necessarily unique) number of genomes
does this amount to?**

```
sum(hpls.gt3[,2:10])  # 80886
```

```
## [1] 80886
```

**… and per sample?**

```
colSums(hpls.gt3[,2:10])
```

```
##   X01   X02   X03   X08   X09   X10   X11   X12   X15 
##  6488  7190  7910 11953 10248  6261  9208  9676 11952
```

```
#  X01   X02   X03   X08   X09   X10   X11   X12   X15 
# 6488  7190  7910 11953 10248  6261  9208  9676 11952 
colSums(hpls.gt3[,11:12]) # protocol 2
```

```
##   X16   X17 
##  9349 16305
```

```
#  X16   X17 
# 9349 16305
```

**How many unique genotypes do we have in the ancestor
population without and with the restriction to >3 reads per
sample?**

```
hpls.ancestor.raw <- hpls[hpls$X01>0,]  # 2622
length(hpls.ancestor.raw[,1])
```

```
## [1] 2622
```

```
hpls.ancestor.gt3 <- hpls.gt3[hpls.gt3$X01>3,]  # 53 haplotypes
length(hpls.ancestor.gt3[,1])
```

```
## [1] 53
```

### Statistics on number of mutations

We define this function that extract all mutations from a population
of genotypes (that potentially carry multiple, partially overlapping
mutations:

```
mutset.pop <- function(hpls=hpls.ancestor.gt3[,1]){
    ms <- NULL
    for(h in hpls){
        ms <- union(strsplit(h,split=";")[[1]], ms) 
    }
    ms
}
```

As an example, the ancestor population, restricted to more than 3
reads, contains the following mutations:

```
mutset.pop(hpls=hpls.ancestor.gt3[,1])
```

```
##  [1] "1303:C->G" "4979:iG"   "1440:iT"   "1306:T->C" "1638:T->G" "538:T->C" 
##  [7] "4401:C->T" "1300:A->G" "1426:G->A" "2447:G->A" "3997:C->T" "2430:iT"  
## [13] "1780:iT"   "859:iA"    "3855:G->T" "472:iT"    "2652:iT"   "58:iA"    
## [19] "4630:C->T" "3339:G->A" "760:C->T"  "5048:G->T" "5005:C->T" "4352:iT"  
## [25] "4575:iT"   "1020:C->T" "1935:C->T" "1726:C->T" "571:T->C"  "2698:C->T"
## [31] "4959:C->T" "2084:C->T" "2645:C->T" "1564:A->G" "607:T->C"  "4588:C->T"
## [37] "927:C->T"
```

**How many unique mutations are in the ancestor
population?**

```
length(mutset.pop(hpls=hpls.ancestor.gt3[,1]))  # 37
```

```
## [1] 37
```

**How many unique mutations, taking together all samples pooled
over every evolutionary line?**

```
length(mutset.pop(hpls=hpls.gt3[,1]))  # 184
```

```
## [1] 184
```

**How many unique mutations do we find in the raw data overall
and in the ancestor?**

```
length(mutset.pop(hpls=hpls[,1]))  # 19758
```

```
## [1] 19758
```

```
length(mutset.pop(hpls=hpls.ancestor.raw[,1]))  # 4582
```

```
## [1] 4582
```

The first number, 19758, is more than 3 times the genome length of
5386. This shows that in the raw data, there are on average multiple
nucleotide substitutions and insertions found at a site.

To look at this in more detail, we define the following
functions:

```
muts.at.pos <- function(pos="2957", h.data=hpls.gt3){
    hpls.mutated <- grep(pos, h.data$haplotype,
                         value=T)
    unique(sapply(hpls.mutated,
                  function(h) strsplit(strsplit(h, pos)[[1]][2],
                                       ";")[[1]][1]))
}

muts.at.pos(pos="2957", h.data=hpls.gt3)
```

```
## [1] ":G->T"
```

```
muts.at.pos(pos="2957", h.data=hpls)
```

```
## [1] ":iA"   ":iG"   ":G->T" ":G->C"
```

**How many new muts evolve compared to the ancestor (as opposed
to new genotypes being recombinants of ancestral ones)?**

```
mutset.2pop.diff <- function(hpls.1=hpls.gt3[hpls.gt3$X02>3,1] , hpls.2=hpls.ancestor.gt3[,1]){
    ms1 <- mutset.pop(hpls.1)
    ms2 <- mutset.pop(hpls.2)
    setdiff(ms1,ms2)
}
head(mutset.2pop.diff(hpls.1=hpls.gt3[,1] , hpls.2=hpls.ancestor.gt3[,1]))
```

```
## [1] "1246:A->G" "1770:G->T" "2969:G->T" "3165:T->C" "3948:iT"   "5257:iA"
```

```
length(mutset.2pop.diff(hpls.1=hpls.gt3[,1] , hpls.2=hpls.ancestor.gt3[,1]))
```

```
## [1] 147
```

This means that out of the 184 mutations we observe in the dataset
(restricted to >3 reads - see above), 147 are not in the ancestor
population and hence must have evolved *de novo*.

### Parallel genotypes

**How many genotypes evolve in parallel in the four independent
evolution lines?** (Constraining the analysis to genotypes that
appear >3.) Hereby a parallel genotype is one that is not present in
the ancestor, but present at later points (generation 196 or 412) in
*multiple* evolution lines. To answer the question, we define the
following functions:

```
is.haplotype.shared <-
    function(h="4630:C->T", h.data=hpls.gt3, verbose=T){
        h.info <- h.data[h.data$haplotype==h,1:10]
        if(dim(h.info)[1]==0)
            if(verbose) stop(paste("Haplotype",h,"does not exist!"))
            else stop()
        else {
            ##check that h not present in ancestor
            in.ancestor <- h.info$X01>3
            if(in.ancestor) {
                shared.across.lines <- FALSE
                if(verbose) warning(paste("Haplotype",h,"found in ancestor population!"))
            }
            else {
                ##presence defined strictly as >3 in each sample
                present.in.line1 <- any(h.info[,c("X02","X10")]>3)
                present.in.line2 <- any(h.info[,c("X03","X11")]>3)
                present.in.line3 <- any(h.info[,c("X08","X12")]>3)
                present.in.line4 <- any(h.info[,c("X09","X15")]>3)
                
                shared.across.lines <-
                    sum(c(present.in.line1,
                          present.in.line2,
                          present.in.line3,
                          present.in.line4))>1                
            }
            return(shared.across.lines)
        }
    }

n.shared.haplotypes <-
    function(h.data=hpls.gt3){ # runs too long with hpls !!!
        sum(sapply(h.data$haplotype, function(h) is.haplotype.shared(h=h, h.data=h.data, verbose=F)))
    }

n.shared.haplotypes(h.data=hpls.gt3)  # 26
```

```
## [1] 26
```

```
#n.shared.haplotypes(h.data=hpls)  # also 26, This runs long!!!
```

**Which are these 26 genotypes?**

```
which.shared.haplotypes <-
    function(h.data=hpls.gt3){ # runs too long with hpls !!!
        h.data[sapply(h.data$haplotype, function(h) is.haplotype.shared(h=h, h.data=h.data, verbose=F)),"haplotype"]
    }

hpls.shared <- which.shared.haplotypes(h.data=hpls.gt3)
hpls.shared
```

```
##  [1] "1300:A->G;1303:C->G;1426:G->A;1638:T->G;1689:C->T;3240:G->C"                              
##  [2] "1303:C->G;1426:G->A;1638:T->G;3165:T->C"                                                  
##  [3] "1300:A->G;1303:C->G;1426:G->A;1638:T->G;1719:C->T;3240:G->C"                              
##  [4] "1300:A->G;1303:C->G;1426:G->A;1638:T->G;1719:C->T;2957:G->T;3240:G->C"                    
##  [5] "1300:A->G;1303:C->G;1426:G->A;1638:T->G;2957:G->T;3240:G->C"                              
##  [6] "1300:A->G;1303:C->G;1426:G->A;1638:T->G;3165:T->C"                                        
##  [7] "1303:C->G;1638:T->G;3165:T->C"                                                            
##  [8] "1300:A->G;1303:C->G;1426:G->A;1638:T->G;1770:G->T;3165:T->C"                              
##  [9] "1300:A->G;1303:C->G;1638:T->G;3165:T->C"                                                  
## [10] "1300:A->G;1303:C->G;1426:G->A;1638:T->G;1734:A->G;3213:T->C"                              
## [11] "1300:A->G;1638:T->G;1734:A->G"                                                            
## [12] "1300:A->G;1638:T->G;2969:G->T"                                                            
## [13] "2969:G->T"                                                                                
## [14] "1300:A->G;2969:G->T"                                                                      
## [15] "1638:T->G;2969:G->T"                                                                      
## [16] "1300:A->G;3165:T->C"                                                                      
## [17] "1300:A->G;1303:C->G;1426:G->A;1638:T->G;1695:C->T;2083:G->A;2969:G->T;3213:T->C;658:C->T" 
## [18] "1300:A->G;1303:C->G;1426:G->A;1638:T->G;1695:C->T;2083:G->A;2969:G->T;3213:T->C"          
## [19] "1300:A->G;1303:C->G;1638:T->G;1695:C->T;2083:G->A;2969:G->T;3213:T->C;658:C->T"           
## [20] "1300:A->G;1303:C->G;1638:T->G;1695:C->T;2083:G->A;2969:G->T;3213:T->C"                    
## [21] "1300:A->G;1638:T->G;1695:C->T;2083:G->A;2969:G->T;3213:T->C"                              
## [22] "1300:A->G;1426:G->A;1638:T->G;1695:C->T;2083:G->A;2969:G->T;3213:T->C;658:C->T"           
## [23] "1300:A->G;1426:G->A;1638:T->G;1695:C->T;2083:G->A;2969:G->T;3213:T->C"                    
## [24] "1300:A->G;1303:C->G;1426:G->A;1638:T->G;1695:C->T;2576:G->T;2969:G->T;3213:T->C"          
## [25] "1300:A->G;1303:C->G;1426:G->A;1638:T->G;1695:C->T;2083:G->A;2969:G->T;3213:T->C;4456:T->C"
## [26] "1300:A->G;1638:T->G;1695:C->T;2083:G->A;2969:G->T;3213:T->C;658:C->T"
```

**Are ancestor genotypes properly excluded?**

```
sum(sapply(hpls.shared, function(h) h %in% hpls.ancestor.gt3$haplotype))
```

```
## [1] 0
```

**How different are the shared haplotypes from the ancestor?
How much have they diverged?** To answer this question, we define
the following functions that calculates the Hamming distance between
genotypes and populations of genotypes:

```
HD <- function(h1,h2){
    ms1 <- strsplit(h1,split=";")[[1]] ## mutset in h1 ([[]] necessary bc strsplit outputs list)
    ms2 <- strsplit(h2,split=";")[[1]] ## mutsset in h2
    ## also possible: length(union(setdiff(ms1,ms2), setdiff(ms2,ms1)))
    length(setdiff(union(ms1,ms2), intersect(ms1,ms2)))
}

## Calculate HD for each shared haplotype from ancestor haplotypes
HD.2pop <- function(hpls.1=hpls.shared, hpls.2=hpls.ancestor.gt3[,1]){
  unname(sapply(hpls.1, function(h1) sapply(hpls.2, function(h2) HD(h1,h2))))
}

## Calculate minimum HD for each shared haplotype from ancestor haplotypes
minHD.2pop <- function(hpls.1=hpls.shared, hpls.2=hpls.ancestor.gt3[,1]){
  unname(sapply(hpls.1, function(h1) min(sapply(hpls.2, function(h2) HD(h1,h2)))))
}

sapply(0:10, function(h) sum(minHD.2pop()==h))
```

```
##  [1]  0 10  5  1  5  5  0  0  0  0  0
```

```
summary(minHD.2pop())
```

```
##    Min. 1st Qu.  Median    Mean 3rd Qu.    Max. 
##   1.000   1.000   2.000   2.615   4.000   5.000
```

**How different are the evolved haplotypes from each other
after 196 vs 412 generations?**

Evolutionary line 1:

```
sapply(0:10, function(h) sum(minHD.2pop(hpls.1=hpls.gt3[hpls.gt3[,"X02"]>0,1], hpls.2=hpls.ancestor.gt3[,1])==h))
```

```
##  [1] 12 23 34 38  0  0  0  0  0  0  0
```

```
sapply(0:10, function(h) sum(minHD.2pop(hpls.1=hpls.gt3[hpls.gt3[,"X10"]>0,1], hpls.2=hpls.ancestor.gt3[,1])==h))
```

```
##  [1]  4 19 25 35 30 23  0  0  0  0  0
```

```
sapply(0:10, function(h) sum(minHD.2pop(hpls.1=hpls.gt3[hpls.gt3[,"X10"]>0,1], hpls.2=hpls.gt3[hpls.gt3[,"X02"]>0,1])==h))
```

```
##  [1] 31 34 32 11 13 14  1  0  0  0  0
```

Evolutionary line 2:

```
sapply(0:10, function(h) sum(minHD.2pop(hpls.1=hpls.gt3[hpls.gt3[,"X03"]>0,1], hpls.2=hpls.ancestor.gt3[,1])==h))
```

```
##  [1] 10 32 58 35  1  0  0  0  0  0  0
```

```
sapply(0:10, function(h) sum(minHD.2pop(hpls.1=hpls.gt3[hpls.gt3[,"X11"]>0,1], hpls.2=hpls.ancestor.gt3[,1])==h))
```

```
##  [1]  3 13 18 40 39 49  2  0  0  0  0
```

```
sapply(0:10, function(h) sum(minHD.2pop(hpls.1=hpls.gt3[hpls.gt3[,"X11"]>0,1], hpls.2=hpls.gt3[hpls.gt3[,"X03"]>0,1])==h))
```

```
##  [1] 46 51 64  3  0  0  0  0  0  0  0
```

Divergence of evolution line 1 versus 2, after generation 196 and
412:

```
sapply(0:10, function(h) sum(minHD.2pop(hpls.1=hpls.gt3[hpls.gt3[,"X02"]>0,1], hpls.2=hpls.gt3[hpls.gt3[,"X03"]>0,1])==h))
```

```
##  [1] 10 24 33 40  0  0  0  0  0  0  0
```

```
sapply(0:10, function(h) sum(minHD.2pop(hpls.1=hpls.gt3[hpls.gt3[,"X10"]>0,1], hpls.2=hpls.gt3[hpls.gt3[,"X11"]>0,1])==h))
```

```
##  [1] 41  8 16 25 19 12 15  0  0  0  0
```

### Parallel mutations

**How many mutations evolve in parallel in the four independent
evolution lines?** (Constraining the analysis to genotypes that
appear >3.) Hereby a parallel mutation is one that is not present in
the ancestor, but present at later points (generation 196 or 412) in
*multiple* evolution lines.

To answer the question, we define the following functions:

```
is.mut.shared <-
    function(mut="1755:A->G", h.data=hpls.gt3, verbose=T){
        h.info <- h.data[grepl(mut, h.data$haplotype),1:10]
        if(dim(h.info)[1]==0)
            if(verbose) stop(paste("Mutation",mut,"not found!"))
            else stop()
        else {
            ##check that mut not present in ancestor
            in.ancestor <- any(h.info$X01>3)
            if(in.ancestor) {
                shared.across.lines <- FALSE
                present.in.line1 <- FALSE
                present.in.line2 <- FALSE
                present.in.line3 <- FALSE
                present.in.line4 <- FALSE
                if(verbose){
                    warning(paste("Mutation",mut,
                                  "found in ancestor population!"))
                }
            }
            else {
                present.in.line1 <- any(h.info[,c("X02","X10")]>3)
                present.in.line2 <- any(h.info[,c("X03","X11")]>3)
                present.in.line3 <- any(h.info[,c("X08","X12")]>3)
                present.in.line4 <- any(h.info[,c("X09","X15")]>3)
                
                shared.across.lines <-
                    sum(c(present.in.line1,
                          present.in.line2,
                          present.in.line3,
                          present.in.line4))>1
            }
            return(
                list(is.shared=shared.across.lines,
                     in.how.many.lines=sum(c(present.in.line1,
                                             present.in.line2,
                                             present.in.line3,
                                             present.in.line4)),
                     in.which.lines=(0:4)[c(in.ancestor,
                                            present.in.line1,
                                            present.in.line2,
                                            present.in.line3,
                                            present.in.line4)])
            )
        }
    }

n.shared.muts <-
    function(h.data=hpls.gt3){ # runs too long with hpls !!!
        sum(sapply(mutset.pop(h.data$haplotype),
                   function(m) is.mut.shared(mut=m,
                                             h.data=h.data,
                                             verbose=F)$is.shared))
    }

n.shared.muts(h.data=hpls.gt3)  # 47
```

```
## [1] 47
```

```
length(mutset.pop(hpls.gt3$haplotype))  # 184
```

```
## [1] 184
```

Thus, out of the 184 mutations in our data set, 47 occur in multiple
evolutionary lines.

**How many mutations evolve in all four evolution
lines?** (Again, constraining the analysis to genotypes that
appear >3.)

```
sum(sapply(mutset.pop(hpls.gt3$haplotype),
           function(m) is.mut.shared(mut=m,
                                     h.data=hpls.gt3,
                                     verbose=F)$in.how.many.lines)==4)
```

```
## [1] 7
```

**How many of the parallel mutations are
synonymous?**

To this end, we define the following helper function that determines
the gene(s) in which a mutation is:

```
in.which.gene <- function(mut="1300:A->G"){
    locus <- as.integer(strsplit(mut,split=":")[[1]][1]) + 1 # because python indexes start with 9
    phiX174.genes <- read.table("../../python-code/data/phiX174_genes.txt", header=T, sep="\t")

    ## get list of genes that contain locus
    ind1 <- ((phiX174.genes$start_position_on_the_genomic_accession <= locus))
    ind2 <- ((phiX174.genes$end_position_on_the_genomic_accession >= locus))
    ind <- ind1 & ind2
    return(phiX174.genes$description[ind])
}
```

The genes depend only on the locus, not the nucleotide
substitution.

With this helper function we can define the following function that
tests if a mutation is synonymous in all of the genes in which it
sits:

```
is.mut.syn <- function(mut="1300:A->G"){
    
    locus <- as.integer(strsplit(mut,split=":")[[1]][1]) + 1 # because python indexes start with 0
    mut.type <- strsplit(mut,split=":")[[1]][2]

    ## this checks if the mut is other than a substitution
    ## (mostly insertions), in which case the mutation is assumed
    ## to be not synonymous  
    if(!grepl("->",mut)){ 
        is.syn <- FALSE
    }
    else {
        nt.orig <- strsplit(mut.type, split="->")[[1]][1]
        nt.subs <- tolower(strsplit(mut.type, split="->")[[1]][2])
    
        require(seqinr)
        ## get ref seq:
        phiX174.ref.seq <- read.fasta(file="../../python-code/data/PhiX174_NCBI.fasta",
                                      seqtype="DNA")[[1]]
        phiX174.consensus <- read.fasta(file="../../python-code/data/consensus_start_run2.fasta",
                                        seqtype="DNA")[[1]]
        ## get starts of ORFs: phiX174_genes.txt
        phiX174.genes <- read.table("../../python-code/data/phiX174_genes.txt", header=T, sep="\t")
        
        ## get list of genes that contain locus
        ind1 <- ((phiX174.genes$start_position_on_the_genomic_accession <= locus))
        ind2 <- ((phiX174.genes$end_position_on_the_genomic_accession >= locus))
        ind <- ind1 & ind2

        if(any(ind)){ #this checks if locus is in any gene
            genes.with.mut <- phiX174.genes$description[ind]
    
            is.syn <- vector("logical", length=length(genes.with.mut))
            names(is.syn) <- genes.with.mut
        
            for (i in length(genes.with.mut)){
                f <- locus - phiX174.genes$start_position_on_the_genomic_accession[ind][i]
                kodon.orig <- phiX174.consensus[0:2 + locus - f%%3]
                kodon.subs <- kodon.orig
                kodon.subs[1+f%%3] <- nt.subs
        
                is.syn[i] <- (c2s(kodon.subs) %in% syncodons(c2s(kodon.orig))[[1]])

            ## check by translating both codons:
            ##aa.orig <- translate(kodon.orig, frame=0)
            ##print(aa.orig)
            ##aa.subs <- translate(kodon.subs, frame=0)
            ##print(aa.subs)
            }
        }
        else is.syn <- TRUE
        ## because if the mutation is not in any gene
        ## it will not affect translated protein.
        ## Technically, it is a mutation/locus in an
        ## "untranslated" region
    }

    return(list(locus=locus,
                mut.type=mut.type,
                is.syn,
                overall=all(is.syn)))
}
```

In this function, we defined a mutation in untranslated regions as
“synonymous”. Strictly speaking, the output `TRUE` means
**not non-synonymous**, rather than synonymous.

Since phiX174 has overlapping reading frames, some mutations can be
synomymous in one and non-sysnomymous in another:

```
is.mut.syn(mut="100:A->C")
```

```
## $locus
## [1] 101
## 
## $mut.type
## [1] "A->C"
## 
## [[3]]
##     A    A*     K 
## FALSE FALSE  TRUE 
## 
## $overall
## [1] FALSE
```

The above function can be wrapped to check all the mutations in a set
of genotypes:

```
how.many.syn.muts <- function(hpls=hpls.ancestor.gt3$haplotype){
    muts <- mutset.pop(hpls)
    n.muts <- length(muts)
    n.muts.syn <- sum(sapply(muts,
                             function(m) is.mut.syn(m)$overall))
    c(number.muts=n.muts,
      number.muts.syn=n.muts.syn)
}
```

Let’s apply this to the genotypes in the entire dataset, the
ancestor, and the genotypes that emerge in multiple evolution lines:

```
how.many.syn.muts(hpls=hpls.gt3$haplotype)
```

```
##     number.muts number.muts.syn 
##             184              46
```

```
how.many.syn.muts(hpls=hpls.ancestor.gt3$haplotype)
```

```
##     number.muts number.muts.syn 
##              37               7
```

```
how.many.syn.muts(hpls=hpls.shared)
```

```
##     number.muts number.muts.syn 
##              18               6
```

This shows that of the 18 mutations in the shared genotypes 6 are
synonymous, i.e. a fraction of 0.33 - a higher fraction than in the
ancestor (7/37=0.19) or in the entire data set (46/184=0.25).

**Table of all 184 mutations**

To look at these questions more systematically, we calculate the
following table.

```
all.muts <- mutset.pop(hpls.gt3$haplotype)
mut.table <-
    data.frame(mut=all.muts,
               in.ancestor=sapply(all.muts,
                                  function(m)
                                      m %in%
                                      mutset.pop(hpls.ancestor.gt3$haplotype)),
               in.how.many.lines=sapply(all.muts,
                                       function(m)
                                           is.mut.shared(mut=m,
                                                         h.data=hpls.gt3,
                                                         verbose=F)$in.how.many.lines),
               is.shared=sapply(all.muts,
                                function(m)
                                    is.mut.shared(mut=m,
                                                  h.data=hpls.gt3,
                                                  verbose=F)$is.shared),
               is.syn=sapply(all.muts, function(m) is.mut.syn(m)$overall),
               row.names = NULL)
mut.table
```

| mut | in.ancestor | in.how.many.lines | is.shared | is.syn |
| --- | --- | --- | --- | --- |
| 1246:A->G | FALSE | 1 | FALSE | FALSE |
| 1300:A->G | TRUE | 0 | FALSE | FALSE |
| 1303:C->G | TRUE | 0 | FALSE | FALSE |
| 1426:G->A | TRUE | 0 | FALSE | FALSE |
| 1638:T->G | TRUE | 0 | FALSE | FALSE |
| 1770:G->T | FALSE | 3 | TRUE | TRUE |
| 2969:G->T | FALSE | 3 | TRUE | FALSE |
| 3165:T->C | FALSE | 3 | TRUE | FALSE |
| 3948:iT | FALSE | 3 | TRUE | FALSE |
| 5257:iA | FALSE | 1 | FALSE | FALSE |
| 3006:iA | FALSE | 1 | FALSE | FALSE |
| 859:iA | TRUE | 0 | FALSE | FALSE |
| 5304:iT | FALSE | 2 | TRUE | FALSE |
| 3139:iT | FALSE | 1 | FALSE | FALSE |
| 4445:iA | FALSE | 1 | FALSE | FALSE |
| 3819:iT | FALSE | 2 | TRUE | FALSE |
| 4924:iT | FALSE | 4 | TRUE | FALSE |
| 3431:iA | FALSE | 2 | TRUE | FALSE |
| 4408:iC | FALSE | 1 | FALSE | FALSE |
| 4759:C->A | FALSE | 1 | FALSE | FALSE |
| 4753:iT | FALSE | 1 | FALSE | FALSE |
| 780:A->G | FALSE | 1 | FALSE | FALSE |
| 1318:G->A | FALSE | 1 | FALSE | FALSE |
| 2696:T->C | FALSE | 1 | FALSE | TRUE |
| 3202:C->A | FALSE | 1 | FALSE | TRUE |
| 1032:G->T | FALSE | 1 | FALSE | FALSE |
| 2502:iG | FALSE | 2 | TRUE | FALSE |
| 4633:T->C | FALSE | 1 | FALSE | FALSE |
| 969:G->A | FALSE | 4 | TRUE | TRUE |
| 4108:C->T | FALSE | 1 | FALSE | TRUE |
| 2332:iT | FALSE | 1 | FALSE | FALSE |
| 4616:iT | FALSE | 2 | TRUE | FALSE |
| 4382:iT | FALSE | 4 | TRUE | FALSE |
| 4344:iA | FALSE | 4 | TRUE | FALSE |
| 1955:iT | FALSE | 4 | TRUE | FALSE |
| 1440:iT | TRUE | 0 | FALSE | FALSE |
| 2933:iT | FALSE | 2 | TRUE | FALSE |
| 2000:iT | FALSE | 2 | TRUE | FALSE |
| 702:iA | FALSE | 2 | TRUE | FALSE |
| 454:G->A | FALSE | 1 | FALSE | TRUE |
| 4121:G->A | FALSE | 1 | FALSE | FALSE |
| 655:C->T | FALSE | 1 | FALSE | FALSE |
| 4979:iG | TRUE | 0 | FALSE | FALSE |
| 1734:A->G | FALSE | 3 | TRUE | TRUE |
| 4400:G->A | FALSE | 1 | FALSE | FALSE |
| 3246:G->T | FALSE | 1 | FALSE | FALSE |
| 526:T->C | FALSE | 1 | FALSE | TRUE |
| 1780:iT | TRUE | 0 | FALSE | FALSE |
| 4167:A->G | FALSE | 1 | FALSE | FALSE |
| 2652:iT | TRUE | 0 | FALSE | FALSE |
| 472:iT | TRUE | 0 | FALSE | FALSE |
| 4803:C->T | FALSE | 1 | FALSE | FALSE |
| 58:iA | TRUE | 0 | FALSE | FALSE |
| 807:iA | FALSE | 4 | TRUE | FALSE |
| 475:iA | FALSE | 3 | TRUE | FALSE |
| 5089:iA | FALSE | 2 | TRUE | FALSE |
| 3974:iT | FALSE | 4 | TRUE | FALSE |
| 3642:C->T | FALSE | 1 | FALSE | FALSE |
| 1701:T->C | FALSE | 2 | TRUE | TRUE |
| 3099:A->G | FALSE | 1 | FALSE | FALSE |
| 2430:iT | TRUE | 0 | FALSE | FALSE |
| 4084:T->C | FALSE | 1 | FALSE | TRUE |
| 4352:iT | TRUE | 0 | FALSE | FALSE |
| 1689:C->T | FALSE | 2 | TRUE | TRUE |
| 2084:C->T | TRUE | 0 | FALSE | FALSE |
| 2957:G->T | FALSE | 2 | TRUE | FALSE |
| 3240:G->C | FALSE | 3 | TRUE | FALSE |
| 1719:C->T | FALSE | 2 | TRUE | TRUE |
| 3454:G->T | FALSE | 1 | FALSE | FALSE |
| 5002:iT | FALSE | 1 | FALSE | FALSE |
| 2124:G->A | FALSE | 1 | FALSE | TRUE |
| 3001:C->T | FALSE | 1 | FALSE | TRUE |
| 3766:iT | FALSE | 1 | FALSE | FALSE |
| 3337:T->C | FALSE | 1 | FALSE | TRUE |
| 4726:iT | FALSE | 2 | TRUE | FALSE |
| 2558:iT | FALSE | 2 | TRUE | FALSE |
| 1905:C->T | FALSE | 1 | FALSE | TRUE |
| 4575:iT | TRUE | 0 | FALSE | FALSE |
| 2485:iT | FALSE | 2 | TRUE | FALSE |
| 1498:iA | FALSE | 3 | TRUE | FALSE |
| 2296:T->C | FALSE | 1 | FALSE | TRUE |
| 2663:T->C | FALSE | 1 | FALSE | TRUE |
| 4726:T->C | FALSE | 1 | FALSE | FALSE |
| 4687:T->C | FALSE | 1 | FALSE | FALSE |
| 3797:G->A | FALSE | 1 | FALSE | FALSE |
| 3560:G->A | FALSE | 2 | TRUE | FALSE |
| 1954:G->A | FALSE | 1 | FALSE | FALSE |
| 1695:C->T | FALSE | 3 | TRUE | TRUE |
| 2083:G->A | FALSE | 3 | TRUE | FALSE |
| 3213:T->C | FALSE | 3 | TRUE | FALSE |
| 4456:T->C | FALSE | 2 | TRUE | TRUE |
| 2576:G->T | FALSE | 2 | TRUE | FALSE |
| 673:T->C | FALSE | 1 | FALSE | FALSE |
| 3388:G->A | FALSE | 1 | FALSE | TRUE |
| 2847:G->A | FALSE | 1 | FALSE | FALSE |
| 1610:A->G | FALSE | 1 | FALSE | FALSE |
| 1278:iT | FALSE | 1 | FALSE | FALSE |
| 1969:T->C | FALSE | 1 | FALSE | TRUE |
| 1308:T->C | FALSE | 1 | FALSE | TRUE |
| 4816:C->T | FALSE | 1 | FALSE | FALSE |
| 3241:A->G | FALSE | 1 | FALSE | TRUE |
| 3134:C->T | FALSE | 1 | FALSE | FALSE |
| 658:C->T | FALSE | 2 | TRUE | FALSE |
| 1536:G->A | FALSE | 1 | FALSE | TRUE |
| 2792:C->T | FALSE | 1 | FALSE | TRUE |
| 4630:C->T | TRUE | 0 | FALSE | FALSE |
| 4837:A->G | FALSE | 1 | FALSE | FALSE |
| 2103:C->T | FALSE | 1 | FALSE | TRUE |
| 1341:C->T | FALSE | 1 | FALSE | TRUE |
| 4735:G->A | FALSE | 1 | FALSE | FALSE |
| 2966:C->T | FALSE | 1 | FALSE | FALSE |
| 2049:G->T | FALSE | 1 | FALSE | FALSE |
| 968:iA | FALSE | 1 | FALSE | FALSE |
| 4771:C->T | FALSE | 1 | FALSE | FALSE |
| 1136:C->T | FALSE | 1 | FALSE | FALSE |
| 1755:A->G | FALSE | 1 | FALSE | TRUE |
| 3119:C->T | FALSE | 2 | TRUE | FALSE |
| 4432:G->T | FALSE | 1 | FALSE | FALSE |
| 1812:T->C | FALSE | 1 | FALSE | TRUE |
| 2313:G->T | FALSE | 1 | FALSE | TRUE |
| 2314:A->C | FALSE | 1 | FALSE | TRUE |
| 2259:iT | FALSE | 1 | FALSE | FALSE |
| 1821:G->T | FALSE | 1 | FALSE | TRUE |
| 1503:C->T | FALSE | 2 | TRUE | TRUE |
| 5332:iT | FALSE | 1 | FALSE | FALSE |
| 4495:iT | FALSE | 3 | TRUE | FALSE |
| 2372:iT | FALSE | 3 | TRUE | FALSE |
| 212:iA | FALSE | 3 | TRUE | FALSE |
| 1675:iA | FALSE | 1 | FALSE | FALSE |
| 1872:iT | FALSE | 2 | TRUE | FALSE |
| 2544:G->T | FALSE | 1 | FALSE | FALSE |
| 2181:iT | FALSE | 1 | FALSE | FALSE |
| 2075:iT | FALSE | 1 | FALSE | FALSE |
| 819:iA | FALSE | 1 | FALSE | FALSE |
| 1346:A->G | FALSE | 2 | TRUE | FALSE |
| 1305:T->C | FALSE | 1 | FALSE | TRUE |
| 2015:A->G | FALSE | 2 | TRUE | FALSE |
| 1345:G->A | FALSE | 1 | FALSE | FALSE |
| 2145:A->G | FALSE | 2 | TRUE | TRUE |
| 2255:iC | FALSE | 1 | FALSE | FALSE |
| 893:iA | FALSE | 1 | FALSE | FALSE |
| 2936:iG | FALSE | 1 | FALSE | FALSE |
| 4609:A->G | FALSE | 1 | FALSE | FALSE |
| 1300:A->T | FALSE | 1 | FALSE | FALSE |
| 3059:A->G | FALSE | 1 | FALSE | FALSE |
| 2897:T->C | FALSE | 1 | FALSE | TRUE |
| 1862:C->T | FALSE | 1 | FALSE | FALSE |
| 3103:C->A | FALSE | 1 | FALSE | TRUE |
| 1295:A->G | FALSE | 1 | FALSE | FALSE |
| 4376:A->G | FALSE | 1 | FALSE | FALSE |
| 3360:T->C | FALSE | 1 | FALSE | FALSE |
| 3687:G->T | FALSE | 1 | FALSE | FALSE |
| 2978:G->A | FALSE | 1 | FALSE | FALSE |
| 3351:C->T | FALSE | 1 | FALSE | FALSE |
| 2858:C->T | FALSE | 1 | FALSE | TRUE |
| 3858:A->G | FALSE | 1 | FALSE | FALSE |
| 2954:A->G | FALSE | 1 | FALSE | FALSE |
| 2158:C->T | FALSE | 1 | FALSE | FALSE |
| 1890:iT | FALSE | 1 | FALSE | FALSE |
| 4441:iC | FALSE | 1 | FALSE | FALSE |
| 3197:G->A | FALSE | 1 | FALSE | FALSE |
| 2963:G->T | FALSE | 1 | FALSE | FALSE |
| 3197:G->C | FALSE | 1 | FALSE | FALSE |
| 1306:T->C | TRUE | 0 | FALSE | FALSE |
| 538:T->C | TRUE | 0 | FALSE | TRUE |
| 4401:C->T | TRUE | 0 | FALSE | FALSE |
| 2447:G->A | TRUE | 0 | FALSE | TRUE |
| 3997:C->T | TRUE | 0 | FALSE | TRUE |
| 3855:G->T | TRUE | 0 | FALSE | FALSE |
| 3339:G->A | TRUE | 0 | FALSE | FALSE |
| 760:C->T | TRUE | 0 | FALSE | FALSE |
| 5048:G->T | TRUE | 0 | FALSE | FALSE |
| 5005:C->T | TRUE | 0 | FALSE | FALSE |
| 1020:C->T | TRUE | 0 | FALSE | TRUE |
| 1935:C->T | TRUE | 0 | FALSE | TRUE |
| 1726:C->T | TRUE | 0 | FALSE | FALSE |
| 571:T->C | TRUE | 0 | FALSE | FALSE |
| 2698:C->T | TRUE | 0 | FALSE | FALSE |
| 4959:C->T | TRUE | 0 | FALSE | FALSE |
| 2645:C->T | TRUE | 0 | FALSE | TRUE |
| 1564:A->G | TRUE | 0 | FALSE | FALSE |
| 607:T->C | TRUE | 0 | FALSE | FALSE |
| 4588:C->T | TRUE | 0 | FALSE | FALSE |
| 927:C->T | TRUE | 0 | FALSE | TRUE |

Let’s write this table out into separate files for convenience of the
reviewers of our paper:

```
library("xlsx")
write.xlsx2(mut.table, file="mutations.xlsx", sheetName = "Sheet1",
            col.names = TRUE, row.names = TRUE, append = FALSE)
write.csv(mut.table, file="mutations.csv", quote=FALSE)
```

From the mutation table, we can calculate that of the 47 mutations
that evolved in parallel in multiple lines, 10 are synonymous. While the
fraction of synonymous mutations is lower among the parallel mutations,
this difference is not significant:

```
muts.shared.vs.syn <-
    matrix(c(10,37,36,101),
           nrow=2,
           dimnames=list(c("synonymous","nonsynonymous"),
                         c("parallel","unique")))
muts.shared.vs.syn
```

```
##               parallel unique
## synonymous          10     36
## nonsynonymous       37    101
```

```
fisher.test(muts.shared.vs.syn)
```

```
## 
##  Fisher's Exact Test for Count Data
## 
## data:  muts.shared.vs.syn
## p-value = 0.5621
## alternative hypothesis: true odds ratio is not equal to 1
## 95 percent confidence interval:
##  0.3047836 1.7613884
## sample estimates:
## odds ratio 
##  0.7593611
```

### Linkage disequilibrium analysis

The following functions calculates the genomic distance of two
mutations and their linkage disequilibrium:

```
distance.on.genome <- function(mut1="1300:A->G",
                               mut2="1303:C->G"){
    abs(as.integer(strsplit(mut1,split=":")[[1]][1])-as.integer(strsplit(mut2,split=":")[[1]][1]))
}

LD <- function(mut1="1300:A->G", ## the two first ancestor pop muts above
               mut2="1303:C->G", ## the two first ancestor pop muts above
               pop=hpls.gt3[,c("haplotype","X01")], ## make sure this contains a meaningful population, eg a single sample in which recomb could have occurred
               LD.measure="r2", ## "D", or "Dprime", or "r2" 
               warnings=F){ 

    ## number of genomes in pop:
    n <- sum(pop[,2])
    #cat(paste("n =",n, "\n"))
    
    f1 <- sum(pop[pop$haplotype %in% grep(mut1, pop$haplotype, value=T),2])/n
    #cat(paste("From pop: f1 =", f1, "\n")) 
    f2 <- sum(pop[pop$haplotype %in% grep(mut2, pop$haplotype, value=T),2])/n
    #cat(paste("From pop: f2 =", f2, "\n")) 
    f12 <- sum(pop[pop$haplotype %in% grep(mut2, grep(mut1, pop$haplotype, value=T), value=T),2])/n
    #cat(paste("From pop: f12 =", f12, "\n")) 

    if(warnings){
        if(f1==0) message("f1=0")
        if(f2==0) message("f2=0")
        if(f12==0) message("f12=0")
    }

    D <- f12 - f1*f2
    
    if(LD.measure=="D") return(D)
    if(LD.measure=="Dprime" & D<0) return(D/min(c(f1*f2,(1-f1)*(1-f2))))
    if(LD.measure=="Dprime" & D>0) return(D/min(c(f1*(1-f2),(1-f1)*f2)))
    if(LD.measure=="Dprime" & D==0) return(0)
    if(LD.measure=="r2") return(D^2/(f1*(1-f1)*f2*(1-f2)))
}
```

This function calculates all linkage disequilibria in a
population:

```
LD.pop <- function(pop=hpls.gt3[,c("haplotype","X01")],
                   LD.measure="D", ## "D", or "Dprime", or "r2" 
                   output.dist=FALSE){
    mts <- mutset.pop(pop[,1])
    mts <- mts[order(as.numeric(unlist(lapply(strsplit(mts, split=":"),`[[`, 1))))]
    nm <- length(mts)
    LDs<- matrix(NA, nrow=nm, ncol=nm,
                 dimnames=list(mts,mts))
    gd <- matrix(NA, nrow=nm, ncol=nm,
                 dimnames=list(mts,mts))
    for(i in 1:(nm-1)){
        for(j in (i+1):nm){
            LDs[i,j] <- LD(mut1=mts[i], mut2=mts[j], pop=pop, LD.measure=LD.measure, warnings=F)
            if(output.dist) gd[i,j] <- distance.on.genome(mut1=mts[i], mut2=mts[j])
        }
    }

    attr(LDs, "barcode") <- substr(names(pop)[[2]], 2,3)

    if(!output.dist) return(LDs)
    else return(list(LDs,gd))
}
```

Let’s calculate `D` for all samples (this takes ages):

```
for(bc in c("01","02","03","08","09","10","11","12","15")){
    ldp <- LD.pop(pop=hpls.gt3[,c("haplotype",paste0("X",bc))],
                  LD.measure="D",
                  output.dist=T)
    assign(paste0("LD.D.pop.X",bc,".dist"),
           ldp)
}
rm(bc,ldp)
```

**Does D for specific muts decline over generations as in Hartl
& Clark p77: `D(g) = D(g=0)*(1-r*d)^g`?** In this
formula, `g` stands for the generation, `d` for
the genomic distance of the two mutations, and `r`
recombination rate.

This function calculates D for two muts in all samples, normalizing
the value to the first timepoint g=0:

```
D.vs.g <- function(mut1="1300:A->G",mut2="1303:C->G",
                   normalize.to.D0=T){
    ##extract LDD for ancestor:
    D0 <- LD.D.pop.X01.dist[[1]][mut1, mut2]
    gd <- LD.D.pop.X01.dist[[2]][mut1, mut2] ##distance.on.genome(mut1, mut2)

    if(normalize.to.D0) normalizer <- D0
    else normalizer <- 1
    D.vs.g.out <- data.frame(mut1=mut1, mut2=mut2,
                             D=D0/normalizer,
                             generation=0, line=0,
                             gen.dist=gd)
    for(bc in c("02","03","08","09","10","11","12","15")){
        D <- get(paste0("LD.D.pop.X",bc,".dist"))[[1]][mut1, mut2]
        l <- barcds[barcds$barcode==bc,4]
        if(barcds[barcds$barcode==bc,3]==1) g <- 196
        if(barcds[barcds$barcode==bc,3]==2) g <- 412
        D.vs.g.out <- rbind(D.vs.g.out,
                            c(mut1,mut2,
                              D/normalizer,
                              ##log(D/normalizer),
                              g,l,gd))
    }
    class(D.vs.g.out$D) <- "numeric"
    class(D.vs.g.out$generation) <- "numeric"
    class(D.vs.g.out$gen.dist) <- "numeric"
    return(D.vs.g.out)
}

## THE FOLLOWING IS NOT JUST FOR X01 (ALTHOUGH THE OTHER SAMPLES ARE NOT MENTIONED IN THE CODE)
## because D.vs.g above calculates the value in each sample!
all.D.vs.g.vs.gd <- function(normalize.to.D0=T){
    output <- NULL
    for(i in 1:(dim(LD.D.pop.X01.dist[[1]])[1]-1)){
        for(j in (i+1):dim(LD.D.pop.X01.dist[[1]])[1]){
            output <- rbind(output,
                            D.vs.g(mut1=dimnames(LD.D.pop.X01.dist[[1]])[[1]][i],
                                   mut2=dimnames(LD.D.pop.X01.dist[[1]])[[1]][j],
                                   normalize.to.D0=normalize.to.D0))
        }
    }
    return(output)
}

D.vs.g.vs.gd.data.0 <- all.D.vs.g.vs.gd()
ind.meaningful <- ( !is.na(D.vs.g.vs.gd.data.0$D)
    & D.vs.g.vs.gd.data.0$D!=Inf
    & D.vs.g.vs.gd.data.0$D!=-Inf )
D.vs.g.vs.gd.data <- D.vs.g.vs.gd.data.0[ind.meaningful,]
    
summary(nls(D~(1-r*gen.dist)^generation,
            data=D.vs.g.vs.gd.data[D.vs.g.vs.gd.data$generation>0,],
            start=list(r=0)))
```

```
## 
## Formula: D ~ (1 - r * gen.dist)^generation
## 
## Parameters:
##    Estimate Std. Error t value Pr(>|t|)
## r 0.0003977  0.0014815   0.268    0.788
## 
## Residual standard error: 133.6 on 5623 degrees of freedom
## 
## Number of iterations to convergence: 9 
## Achieved convergence tolerance: 7.638e-06
```

Thus, the decline of `D` over generation and genomic
distance is not significantly different from zero.

### Testing the consistency of the dsDNA and the ssDNA extraction protocols

We used the “new” dsDNA extraction protocol for all samples from the
four lines and the two time points. In addition to the dsDNA extraction
protocol, we applied the ssDNA extraction to two samples. These have the
barcodes 16 and 17, and correspond to the dsDNA extraction protocol
samples with barcodes 08 and 15.

The following code produces Figure S1A:

```
h.counts.X16.old <- hpls.gt3[hpls.gt3$X08>3 | hpls.gt3$X16>3 ,"X16"]
h.counts.X08.new <- hpls.gt3[hpls.gt3$X08>3 | hpls.gt3$X16>3 ,"X08"]
h.counts.X17.old <- hpls.gt3[hpls.gt3$X17>3 | hpls.gt3$X15>3 ,"X17"]
h.counts.X15.new <- hpls.gt3[hpls.gt3$X17>3 | hpls.gt3$X15>3 ,"X15"]
pdf("../figures/protocol-consistency-corr.pdf", width=9, height=5)
par(mfrow=c(1,2))->op
plot(x=replace(h.counts.X08.new, h.counts.X08.new==0, 0.1),
     y=replace(h.counts.X16.old, h.counts.X16.old==0, 0.1),
     main="line=3, generation=196",
     xlab="", #"Counts of genotypes determined \n by dsDNA extraction protocol",
     ylab="", #"Counts of genotypes determined \n by alternative, ssDNA extraction protocol",
     type="p", pch=16, cex=1, col=rgb(0,0,0,alpha=0.5),
     log="xy", axes=F)
axis(1, at=c(0.1,1,10,100,1000), labels=c("0", "1", "10", "100", "1000"))
axis(2, at=c(0.1,1,10,100,1000), labels=c("0", "1", "10", "100", "1000"))
plot(x=replace(h.counts.X15.new, h.counts.X15.new==0, 0.1),
     y=replace(h.counts.X17.old, h.counts.X17.old==0, 0.1),
     main="line=4, generation=412",
     xlab="", #"Counts of genotypes determined \n by dsDNA extraction protocol",
     ylab="", #"Counts of genotypes determined \n by alternative, ssDNA extraction protocol",
     type="p", pch=16, cex=1, col=rgb(0,0,0,alpha=0.5),
     log="xy", axes=F)
axis(1, at=c(0.1,1,10,100,1000), labels=c("0", "1", "10", "100", "1000"))
axis(2, at=c(0.1,1,10,100,1000), labels=c("0", "1", "10", "100", "1000"))
mtext(text="Counts of genotypes determined by dsDNA extraction protocol",
      side=1,outer=T, line=-1.5)
mtext(text="Counts of genotypes determined \n by alternative, ssDNA extraction protocol",
      side=2,outer=T, line=-2)
par(op); rm(op)
dev.off()
```

```
## png 
##   2
```

The counts of the genotypes generated by the two protocols are
significantly correlated:

```
cor.test(h.counts.X15.new, h.counts.X17.old, method="kendall")
```

```
## 
##  Kendall's rank correlation tau
## 
## data:  h.counts.X15.new and h.counts.X17.old
## z = 7.8898, p-value = 3.026e-15
## alternative hypothesis: true tau is not equal to 0
## sample estimates:
##       tau 
## 0.4360455
```

```
cor.test(h.counts.X08.new, h.counts.X16.old, method="kendall")
```

```
## 
##  Kendall's rank correlation tau
## 
## data:  h.counts.X08.new and h.counts.X16.old
## z = 9.1751, p-value < 2.2e-16
## alternative hypothesis: true tau is not equal to 0
## sample estimates:
##       tau 
## 0.5707623
```

The following code produces Figure S1B:

```
pdf("../figures/protocol-consistency-count-distro.pdf", width=9, height=9)
par(mfcol=c(2,2), xpd=T)->op
plot(h.counts.X08.new[order(h.counts.X08.new, decreasing=T)],
     main="line=3, generation=196",
     type="h",log="y", col=rgb(0,0,0,alpha=0.5),
     axes=F, xlab="", ylab="")
```

```
## Warning in xy.coords(x, y, xlabel, ylabel, log): 1 y value <= 0 omitted from
## logarithmic plot
```

```
lines(h.counts.X16.old[order(h.counts.X08.new, decreasing=T)],
      type="h", col=rgb(1,0,0,alpha=0.5))
#replace(h.counts.X15.new, h.counts.X15.new==0, 0.1),
axis(1)
axis(2, at=c(1,10,100,1000), labels=c("1", "10", "100", "1000"))
mtext(text="Genotype index (ordered by decreasing counts according to dsDNA extraction protocol)",
      side=1, line=3.5, adj=-0.2)

plot(h.counts.X16.old[order(h.counts.X16.old, decreasing=T)],
     #main="line=3, generation=196",
     type="h",log="y", col=rgb(1,0,0,alpha=0.5),
     axes=F, xlab="", ylab="")
```

```
## Warning in xy.coords(x, y, xlabel, ylabel, log): 30 y values <= 0 omitted from
## logarithmic plot
```

```
lines(h.counts.X08.new[order(h.counts.X16.old, decreasing=T)],
      type="h", col=rgb(0,0,0,alpha=0.5))
#replace(h.counts.X15.new, h.counts.X15.new==0, 0.1),
axis(1)
axis(2, at=c(1,10,100,1000), labels=c("1", "10", "100", "1000"))

plot(h.counts.X15.new[order(h.counts.X15.new, decreasing=T)],
     main="line=4, generation=412",
     type="h",log="y", col=rgb(0,0,0,alpha=0.5),
     axes=F, xlab="", ylab="")
```

```
## Warning in xy.coords(x, y, xlabel, ylabel, log): 1 y value <= 0 omitted from
## logarithmic plot
```

```
lines(h.counts.X17.old[order(h.counts.X15.new, decreasing=T)],
      type="h", col=rgb(1,0,0,alpha=0.5))
legend(120,1200,
       legend=c("dsDNA protocol", "ssDNA protocol"),
       fill=c(rgb(0,0,0,alpha=0.5),rgb(1,0,0,alpha=0.5)),
       bty="n")
#replace(h.counts.X15.new, h.counts.X15.new==0, 0.1),
axis(1)
axis(2, at=c(1,10,100,1000), labels=c("1", "10", "100", "1000"))

plot(h.counts.X17.old[order(h.counts.X17.old, decreasing=T)],
     #main="line=3, generation=196",
     type="h",log="y", col=rgb(1,0,0,alpha=0.5),
     axes=F, xlab="", ylab="")
```

```
## Warning in xy.coords(x, y, xlabel, ylabel, log): 21 y values <= 0 omitted from
## logarithmic plot
```

```
lines(h.counts.X15.new[order(h.counts.X17.old, decreasing=T)],
      type="h", col=rgb(0,0,0,alpha=0.5))
#replace(h.counts.X15.new, h.counts.X15.new==0, 0.1),
axis(1)
axis(2, at=c(1,10,100,1000), labels=c("1", "10", "100", "1000"))

mtext(text="Genotype index (ordered by decreasing counts according to ssDNA extraction protocol)",
      side=1,outer=T, line=-1.5)
mtext(text="Counts of genotypes",
      side=2,outer=T, line=-2)
par(op); rm(op)
dev.off()
```

```
## png 
##   2
```

The “new” dsDNA extraction protocol generated is more efficient as it
identifies higher numbers of unique genotypes.

Line 3, generation 196:

```
sum(h.counts.X08.new>3)
```

```
## [1] 129
```

```
sum(h.counts.X16.old>3)
```

```
## [1] 74
```

Line 4, generation 412:

```
sum(h.counts.X15.new>3)
```

```
## [1] 156
```

```
sum(h.counts.X17.old>3)
```

```
## [1] 106
```
